# Mammary Tissue-Derived Extracellular Matrix Hydrogels Reveal the Role of Irradiation in Driving a Pro-Tumor and Immunosuppressive Microenvironment

**DOI:** 10.1101/2022.05.16.492117

**Authors:** Tian Zhu, Steven M. Alves, Arianna Adamo, Xiaona Wen, Kevin C. Corn, Anastasia Shostak, Shereena Johnson, Nicholas D. Shaub, Shannon E. Martello, Benjamin C. Hacker, Antonio D’Amore, Rizia Bardhan, Marjan Rafat

## Abstract

Radiation therapy (RT) is essential for triple negative breast cancer (TNBC) treatment. However, patients with TNBC continue to experience recurrence after RT. The role of the extracellular matrix (ECM) of irradiated breast tissue in tumor recurrence is still unknown. In this study, we evaluated the structure, molecular composition, and mechanical properties of irradiated murine mammary fat pads (MFPs) and developed ECM hydrogels from decellularized tissues (dECM) to assess the effects of RT-induced ECM changes on breast cancer cell behavior. Irradiated MFPs were characterized by increased ECM deposition and fiber density compared to unirradiated controls, which may provide a platform for cell invasion and proliferation. ECM component changes in collagens I, IV, and VI, and fibronectin were observed following irradiation in both MFPs and dECM hydrogels. Encapsulated TNBC cell proliferation and invasive capacity was enhanced in irradiated dECM hydrogels. In addition, TNBC cells co-cultured with macrophages in irradiated dECM hydrogels induced M2 macrophage polarization and exhibited further increases in proliferation. Our study establishes that the ECM in radiation-damaged sites promotes TNBC invasion and proliferation as well as an immunosuppressive microenvironment. This work represents an important step toward elucidating how changes in the ECM after RT contribute to breast cancer recurrence.

## 1. Introduction

Breast cancer is the second leading cause of cancer related mortality in women [1]. One widely used treatment for patients with breast cancer is radiation therapy (RT), which typically enhances antitumor effects and eliminates residual tumor cells after surgery and chemotherapy. However, approximately 20% of breast cancer patients suffer locoregional recurrence after initial treatment, especially in triple negative breast cancer (TNBC) where patients have higher mortality compared to other subtypes [2,3]. Over 13% of treated TNBC patients suffer local recurrence at the primary site [4]. Previous work has shown that radiation of normal tissues under immunocompromised conditions can induce excess immunosuppressive M2 macrophage infiltration, which ultimately leads to tumor cell recruitment, suggesting that recurrence following RT may be dependent on immune status [5–7]. Therefore, understanding the mechanisms behind recurrence post-RT in TNBC is critical for improved patient survival.

The extracellular matrix (ECM) in the tumor microenvironment plays a major role in tumor progression and metastasis. ECM structure, composition, and mechanical properties have profound effects on cell and tissue phenotype [8]. Recent studies have revealed that ECM deposition is altered by RT-induced fibrosis and that tumor cells grow differently on ECM depending on the tissue origin [9,10]. Excess ECM deposition can act as a barrier to shield cancer cells from treatment and negatively affects the transport of oxygen, nutrients, and metabolites, leading to drug resistance and reducing overall survival [11]. We are therefore motivated to determine the effect of the irradiated ECM on tumor and immune cell behavior, which has not yet been explored. A variety of synthetic hydrogels have been designed to study the impact of ECM on cell behavior. However, these synthetic systems have many limitations in recapitulating the native ECM. Decellularized ECM (dECM) hydrogels prepared from mammalian organs retain the composition of the original tissue and are considered promising biological scaffolds for cell growth and tissue regeneration. For example, dECM hydrogels have been engineered from multiple organs, including adipose tissue, dermis, or bladder, and then applied to cell culture *in vitro* or as natural injectable materials to repair and reconstruct tissues *in vivo* [12–14]. In this study, we developed dECM hydrogels to replicate the *in vivo* adipose tissue microenvironment and study the influence of the irradiated ECM on tumor recurrence *in vitro*. This model allows for the analysis of the direct effects of radiation on changes in the ECM within mammary tissue, providing novel insights into strategies for preventing cancer recurrence post-therapy.

We hypothesized that RT-induced ECM alterations contribute to local recurrence by facilitating a pro-tumor microenvironment that encourages tumor cell growth and invasion. To test this hypothesis, we first analyzed the structure, composition, and mechanical properties of murine mammary fat pads (MFPs) following irradiation. We then fabricated dECM hydrogels from MFPs and evaluated the role of the irradiated ECM on tumor and immune cell behavior. This work represents a crucial step toward elucidating how modulation of the ECM after RT contributes to breast cancer recurrence.

## 2. Materials and Methods

### 2.1 Animals

Animal studies were performed in accordance with institutional guidelines and protocols approved by the Vanderbilt University Institutional Animal Care and Use Committee. 8-10-week-old female Nu/Nu mice were obtained from Charles River Laboratories. The mice were allowed free access to standard diet and water and maintained under a 12 hr light/12 hr dark cycle.

### 2.2 Preparation of MFP-derived dECM hydrogels

MFP-derived dECM was prepared as previously described [15]. In brief, MFPs were harvested from sacrificed Nu/Nu mice using CO_2_ asphyxiation followed by cervical dislocation. Tissues were then irradiated to a dose of 20 Gy *ex vivo* using a cesium source to establish the direct effect of radiation on ECM changes in mammary tissue. MFPs were cultured in complete RPMI media at 37 °C/5% CO_2_ incubator for two days and subsequently stored at −80 °C. Thawed MFPs were treated with 0.02% w/v trypsin/0.05% w/v EDTA (Gibco) in deionized water for 1 hr, 3% v/v Triton-X-100 (Sigma-Aldrich) for 1hr, 4% w/v deoxycholic acid (Frontier Scientific) for 1 hr, 1% v/v penicillin-streptomycin (Gibco) at 4 °C overnight, and 4% v/v ethanol/0.1% v/v peracetic acid (Sigma-Aldrich) for 2 hr. MFPs were then treated to 4x 15 min washes of phosphate-buffered saline (PBS) (Quality Biological, Inc.), 100% n-propanol (Fisher Scientific) incubation for 1hr, and 4x 15 min washes of deionized water. Decellularized and delipidated MFPs were then frozen and lyophilized for use in hydrogel preparation. The lyophilized MFP ECM samples were ground into a powder and mixed into 0.1 M hydrochloric acid–pepsin solution (1 mg/mL) (Sigma-Aldrich) at a concentration of 20 mg/mL, and enzymatically digested for 48 hr under a constant stir rate at room temperature. The digestion was ended by titration to pH 7.4 with 1 M NaOH and a 10x PBS solution, bringing the pre-gel solution to 1x PBS. The dECM hydrogel is liquid at room temperature and forms a gel after 30 min at 37 °C.

### 2.3 Cell lines

GFP- and luciferase-labeled 4T1 mouse mammary carcinoma cells were obtained from Dr. Laura Bronsart (Stanford University) in December 2017. Unlabeled 4T1 cells were obtained from ATCC. MDA-MB-231 human breast cancer parental cells were obtained from Dr. Amato Giaccia (Stanford University) in August 2011. MDA-MB-231 cells were transduced with retrovirus particles encoding for the expression of firefly luciferase gene. All cells were cultured at 37°C and 5% CO_2_. Bone marrow derived macrophages (BMDMs) were isolated from the femurs of 8-10-week-old female Nu/Nu mice [16,17]. All cell lines tested negative for *Mycoplasma* contamination with the MycoAlert Mycoplasma detection kit (Lonza). 4T1 cells were cultured in in RPMI-1640 (Gibco), MDA-MB-231 cells were cultured in DMEM (Gibco), and both were supplemented with 10% fetal bovine serum (FBS) (Sigma-Aldrich) and antibiotics (100 U/mL penicillin and 100 mg/mL streptomycin) (Sigma-Aldrich). Isolated BMDMs were cultured in IMDM (Gibco) supplemented with 10% fetal bovine serum (FBS; Sigma-Aldrich), 10 ng/mL macrophage colony stimulating factor (M-CSF; Gibco), and antibiotics (100 U/mL penicillin and 100 mg/mL streptomycin) for 7 days for maturation into macrophages.

### 2.4 Cell culture in dECM hydrogels

GFP- and luciferase-labeled 4T1 or unlabeled 4T1 cells were seeded in dECM hydrogels at a concentration of 1×10^5^ or 5×10^5^ cells/mL of pre-gel solution, respectively. GFP- and luciferase-labeled MDA-MB-231 cells were encapsulated in dECM hydrogels at a concentration of 5×10^5^ or 10×10^5^ cells/mL of pre-gel solution. 100 µL of the cell-gel solution was added into each well of a 16-well chamber slide (Thermo Fisher Scientific). Cells were thoroughly mixed in the pre-gel solution and then incubated at 37°C for 30 min. 100 µL of complete RPMI media for 4T1 cells or complete DMEM media for MDA-MB-231 cells was added to each well. After culturing at 37 °C for 48 hr, cell proliferation was measured by adding luciferin (0.167 mg/mL) to the wells for 10 min and performing bioluminescence imaging (Caliper LifeSciences IVIS Lumina Series III). The GFP-labeled cells were also visualized using fluorescence microscopy (Leica DMi8) at 0, 24, and 48 hr after gelation.

BMDMs were seeded in dECM hydrogels at concentrations of 5×10^5^ or 10×10^5^ cells/mL of pre-gel solution. 100 µL of cell-gel solution was added into each well of a 16-well chamber slide. Cells were thoroughly mixed in the pre-gel solution and then incubated at 37 °C for 30 min. 100 µL of complete IMDM media was added to each well, and cells were cultured at 37°C for 48 hr.

For co-culture experiments, BMDMs (10×10^5^ cells/mL) were stained with CellTrace™ Far Red Cell Proliferation Kit (Thermo Fisher Scientific) immediately before culturing in 100 µL pre-gel solution with GFP- and luciferase-labeled 4T1 cells (5×10^5^ cells/mL). 50 µL of complete IMDM media, 50 µL of complete RPMI (Gibco) media, and 10 ng/mL M-CSF were added to each well after gel formation and subsequently incubated at 37 °C for 48 hr. Both cell lines were visualized using fluorescence microscopy at 0, 24, and 48 hr post-gelation. Proliferation of 4T1 cells was quantified by bioluminescence measurements at 48 hr. Cell number per field for 4T1 cells and BMDMs was quantified using Fiji software (National Institutes of Health) [18].

### 2.5 Live/Dead assay

Cell viability was examined by the LIVE/DEAD^TM^ Cell Viability Assay Kit (Molecular Probes, Inc.) 48 hr after gelation. In brief, encapsulated unlabeled 4T1 cells or BMDMs were washed twice in Dulbecco’s PBS (DPBS) (Gibco). 100 µL of DPBS containing 1 µM calcein AM (excitation/emission = 494/517 nm) and 2 µM ethidium homodimer (excitation/emission = 528/645 nm) was then added to the wells and measured using a Varioskan plate reader (Thermo Fisher Scientific) after 30 min. Cells were then visualized by fluorescence microscopy (Leica DMi8).

### 2.6 Scanning electron microscopy (SEM) tissue preparation and quantification

MFP structure was examined using SEM (Quanta 250 E-SEM), and fiber network characteristics were quantified using a previously developed image analysis algorithm run on MATLAB software [19]. MFPs were fixed in cold 2% v/v glutaraldehyde with 4% paraformaldehyde in 0.1 M sodium cacodylate buffer (Electron Microscopy Sciences) for 4 hr followed by one wash in the same buffer. Fixed tissues were placed in 1% aqueous osmium tetroxide solution (OsO_4_) (Electron Microscopy Sciences) for 1 hr and then washed twice with milli-Q water. MFPs were dehydrated in a graded series of alcohol (50, 70, 90, 100% ethanol in deionized water) for 10 min per wash and then left in 100% ethanol (Thermo Fisher Scientific) overnight at 4°C. After three additional 45 min changes in 100% ethanol, MFPs were critical point dried (Tousimis). After drying, tissues were sputter coated (Ted Pella, Inc.) with a 4.5 nm thick gold/palladium alloy coating and imaged with SEM. A complete set of fiber network descriptors was collected from SEM images of each MFP: fiber alignment, node density (number of fiber intersections per mm^2^), and fiber diameter. Fiber alignment was described through the normalized orientation index where 0% represents a randomly organized (isotropic) network while 100% represents a completely aligned (anisotropic) network. Porosity was described through the mean of the pore size histogram (mm^2^). Automated extraction of these fiber architectural features was achieved with an algorithm, which has been previously described in detail [19]. Briefly, the SEM image is digitally processed by a cascade of steps, including equalization with a 3×3 median filter, local thresholding through the Otsu method, thinning, smoothing, application of morphological operators, skeletonization, binary filtering for Delaunay network refinement, and ultimately the detection of fiber network architecture and its descriptors.

### 2.7 Raman spectroscopy

MFPs were fixed in 10% neutral buffered formalin (NBF) for 24 hr at 4 °C and submerged in 30% sucrose (Sigma-Aldrich) for 48 hr at 4 °C. The fixed samples were then embedded in OCT and cut into 5 µm sections onto Raman grade CaF_2_ disks (Sigma-Aldrich). The Raman measurements were taken with 5 s exposure time using a Renishaw inVia Raman microscope system with a 785 nm laser that delivered ∼30 mW of power. A 100× objective lens was used to focus a laser spot on the surface of samples. Raman spectra were analyzed by a custom MATLAB (R2019b) code to perform smoothing and biological fluorescent background subtraction. The spectra were first smoothed by using the Savitzky and Golay filter with fifth order and coefficient value of 15. Modified polynomial fit method was then performed to remove the background fluorescence by using a 7th order polynomial with a threshold of 0.0001. Afterward, Raman spectra were normalized using min-max normalization method. Principal components analysis (PCA) was performed by using the MATLAB built-in “PCA” function. Two principal components were plotted to discriminate groups.

### 2.8 Atomic force microscopy (AFM)

MFPs or dECM hydrogels were cut into 1mm x 1mm x 1mm pieces, and each specimen was immobilized on 22 mm x 22 mm coverslip with a thin layer of two-component fast drying epoxy glue (Devcon). All AFM indentations were performed using a Bruker inverted optical AFM and silicon nitride triangle cantilever with a 10 μm diameter borosilicate spherical tip (205 mm-long DNP-S10 triangular silicon nitride cantilevers, resonance frequency (air) f =18 kHz, nominal cantilever spring constant *k* = 0.08 Nm, tip radius = 5 µm, Bruker). The exact spring constant (k) of the cantilever was determined before each experiment using the thermal tune method and the deflection sensitivity was determined in fluid using glass substrates as an infinitely stiff reference material. Force-distance (FD) measurements of excised tissues were performed in a fluid cell. Samples were indented with a calibrated force of 5 nN, and the Hertz model for spherical indenters was used to determine the elastic modulus of the tissues.

### 2.9 Luminex multiplex immunoassay

Conditioned media (CM) from GFP- and luciferase-labeled 4T1 cells cultured in dECM hydrogels was collected after 2 days incubation following centrifugation at 1,100 rpm to remove cells. CM was filtered through a 0.2μm PVDF filter (Whatman) and then stored at −80 °C until processed. Three replicates were collected independently. All samples were evaluated using a mouse 32-plex Affymetrix kit (Eve Technologies Corporation). **Supplementary Table S1** lists all cytokines analyzed.

### 2.10 Histological analysis and immunofluorescence (IF)

MFPs were removed from 8-10-week-old female Nu/Nu mice and irradiated to a dose of 20 Gy *ex vivo* using a cesium source. After 48 hr incubation at 37 °C in complete RPMI media, control and irradiated MFPs were fixed in 10% NBF for 48 hr, paraffin-embedded, and sectioned (5 μm). For hematoxylin and eosin (H&E) staining (Sigma-Aldrich), standard procedures were followed, including deparaffinization, hydration, staining with hematoxylin, and counterstaining with eosin. For Masson’s trichrome staining (Thermo Fisher Scientific), sample preparation included deparaffinization, rehydration, staining with Weigert’s iron hematoxylin and Biebrich scarlet-acid fuchsin, differentiation in phosphomolybdic/phosphotungstic acid solution, and staining in aniline blue solution. Collagen area per field was quantified using Fiji. For IF staining, tissue sections (5 μm) were deparaffinized followed by antigen retrieval using citrate buffer (10mM, pH 6). After blocking in 10% goat serum, sections were incubated with primary antibodies, including anti-collagen I (1:200, Thermo Fisher Scientific), anti-collagen IV (1:200, Thermo Fisher Scientific), anti-collagen VI (1:200, Abcam) and anti-fibronectin (1:400, Thermo Fisher Scientific) overnight at 4°C. Fluorescently labeled secondary antibodies (Alexa Fluor 488) were used to stain tissues for 1 hr at room temperature followed by applying Sudan Black B (Acros Organics) to reduce autofluorescence. Sections were mounted using ProLong glass antifade mountant with NucBlue stain (Thermo Fisher Scientific) and imaged using a fluorescence microscope. For F-actin staining, cells after 48 hr incubation in dECM hydrogels were fixed in 10% NBF for 10 min and washed with PBS. Cell membranes were permeabilized by 0.1% Triton X-100 in PBS for 5 min and washed with PBS. Cells were then incubated with phalloidin (Phalloidin-iFluor 594, Abcam) for 1 hr at room temperature and washed with PBS. For cortactin staining, cells after fixation and permeabilization by similar methods were blocked with 5% NGS in PBS for 1 hr at room temperature. Cells were incubated with anti-cortactin (1:50, Novus Biologicals) primary antibody in 1% BSA/PBS overnight at 4 °C followed by three washes in PBS. Cells were then incubated with secondary antibodies (goat anti-rabbit IgG Alexa-Fluor 488; Thermo Fisher Scientific) for 1 hr at RT and washed with PBS. For Ki67 staining, 4T1s and BMDMs encapsulated in dECM hydrogels were fixed, permeabilized, and blocked followed by incubation with anti-Ki67 (1:200, Abcam) primary antibody in 1% BSA/PBS overnight at 4 °C followed by three washes in PBS. Encapsulated cells were then incubated with secondary antibodies (goat anti-rabbit IgG Alexa-Fluor 488 or 594; Thermo Fisher Scientific) for 1 hr at room temperature and washed with PBS. Chambers were removed and mounted with ProLong glass antifade mounting media with NucBlue stain. Samples were imaged using fluorescence microscopy (Leica DMi8), and fluorescence intensity per field was quantified using Raw Integrated Density (RawIntDen) in Fiji. F-actin and cortactin colocalization was determined by identifying the fraction of overlapping pixels for each fluorophore using MATLAB.

### 2.11 Rheology

500 µL pre-gel solution was added onto a rheometer plate (AR 2000ex Rheometer), followed by lowering the arm (25mm) to ensure complete filling of the gap between the plate and the arm with pre-gel solution. A time sweep analysis of the pre-gel solution was conducted for a duration for 10 min following a 30 min incubation at 37 °C with 0.5% applied strain and strain frequency of 0.1 Hz. The Young’s modulus (E) was determined using the following equation:

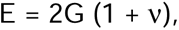

where ν is the Poisson’s ratio (0.5), and G is the bulk modulus determined by the sum of the storage (G’) and loss (G”) modulus [20].

### 2.12 Invasion Assay

A volume of 100 µL of both control and irradiated dECM hydrogels was individually introduced into transwell inserts (Corning, 0.4 µm pore size) and subsequently incubated at 37 °C for 30 min. 750 µL of complete RPMI media was then added into the bottom chamber, and 10^5^ 4T1 cells, suspended in 0.1% BSA in RPMI, were seeded onto either coated invasion or uncoated control migration (8 µm pore) inserts. After a 24 hr incubation period, cells that traversed the inserts or migrated through uncoated inserts were fixed in methanol, and inserts were mounted using Prolong glass antifade mounting media with NucBlue. Fluorescence images were taken, and the number of nuclei per field was quantified. The extent of invasion was determined by dividing the number of invading cells by the number of migrating cells.

### 2.13 Proliferation Assay

5,000 or 15,000 BMDMs were seeded into the dECM hydrogels at 37 °C for 44 hr, and alamarBlue Cell Viability Reagent (Thermo Fisher Scientific) was added directly into each well. Plates were incubated at 37 °C for an additional 4 hr to allow cells to convert resazurin to resorufin to determine proliferation. Subsequentially, fluorescence (excitation: 540nm and emission: 585nm) was measured by a Varioskan plate reader.

### 2.14 Flow Cytometry

Cells were extracted from dECM hydrogels using a 2 mg/mL solution of collagenase II (Sigma) for 30 min at 37 °C. Subsequently, the cells were stained with the fixable Aqua stain (Thermo Fisher Scientific). Simultaneously, FC receptors were blocked with CD16/32 (Biolegend), and cells were stained with surface markers for a duration of 20 mins at 4°C. The panel of surface markers included F4/80 (PE, Biolegend), CD64 (PerCP-eFluor^TM^ 710, Thermo Fisher Scientific), CD86 (eFluor^TM^450 eBioscience^TM^, Thermo Fisher Scientific), IL4Rα (PE/Cyanine7, Biolegend), and integrin β3 (APC, eBioscienceTM, Thermo Fisher Scientific). Following staining, the cells were rinsed with PBS and fixed with 10% NBF for 20 mins at 4°C. Intracellular staining was carried out using an intracellular permeabilization buffer (Thermo Fisher Scientific). Fixed cells underwent a 5 min rinse with PBS, followed by a 5 min rinse with permeabilization buffer. Subsequently, they were incubated with antibodies diluted in permeabilization buffer for a duration of 30 mins at room temperature in the dark. Post-incubation, cells were rinsed with permeabilization buffer and resuspended in PBS. The intracellular marker of interest was CD206 (Alexa-Fluor 488, Biolegend). Flow cytometry was conducted on a four-laser Amnis CellStream machine (Luminex), and FlowJo software was used for data analysis. Compensation was achieved through the utilization of compensation beads (Thermo Fisher Scientific). The following antibody clones were used for analysis: F4/80 (BM8), CD64 (X54-5/7.1), CD86 (GL1), CD206 (C068C2), IL-4Rα (I015F8), Integrin β3 (2C9.G3).

### 2.15 Patient data analysis

We used The Cancer Genome Atlas (TCGA) to obtain patient mRNA expression. The TCGA TARGET GTEx study was accessed through the University of California Santa Cruz’s Xena platform (https://xenabrowser.net/) on May 3^rd^, 2022 to evaluate how gene expression of *Col4A3BP* and *Col4A2-AS1* in the normal adjacent tissue of breast cancer patients correlated with patient outcome [21]. The Xena visualization tool was used to generate Kaplan-Meier curves (https://xenabrowser.net/, n = 113 patients). To investigate breast cancer patient gene expression, we accessed mRNA data from the TCGA cBioPortal (PanCancer dataset, n = 1,082 patients) on May 10^th^, 2022 [22]. From these data, correlations between collagen, invadopodia, and secreted cytokine mRNA expression were determined. We also interrogated publicly available datasets though KMplotter (kmplot.com), which is a meta-analysis tool. Breast cancer patients were filtered based on ER, PR, and HER2 positivity. TNBC patients (n= 126 patients) were used for analysis and stratified based on high or low expression of C-C motif chemokine ligand (*CCL2*) with data reported as probability of survival.

### 2.16 Statistical analysis

In each experiment, obtained data are presented as the mean with standard deviation (SD). Differences among more than two groups were tested using one-way analysis of variance (ANOVA), and Student’s t-tests were performed to compare differences between two groups. p<0.05 was considered significant. All analyses were performed using GraphPad Prism 9.

## 3. Results

### 3.1 Structure and composition of control and irradiated MFPs

SEM images of control and irradiated MFPs revealed more ECM fibers covering irradiated adipose cells than unirradiated adipose cells as well as thinner and denser fibers after radiation (**Fig. 1A-D**). We subsequently quantified ECM fiber network characteristics using fiber network analysis software to confirm our qualitative observations [19] (**Fig.1E-H**). Collagen deposition was evaluated by the fiber covered area ratio parameter, which increased significantly after radiation (p<0.01) (**Fig. 1E**). Visual inspection of the algorithm output showed accurate automatic detection of the fiber network for MFPs (**Supplementary Fig. S1A**). Both fiber diameter and pore size decreased significantly in irradiated MFPs (p<0.05) (**Fig. 1F-G**). The node density of irradiated MFPs also increased significantly (**Fig. 1H**). However, the porosity and orientation index showed no significant changes between control and irradiated samples (**Supplementary Fig. S1B-C**).

**Figure 1.**
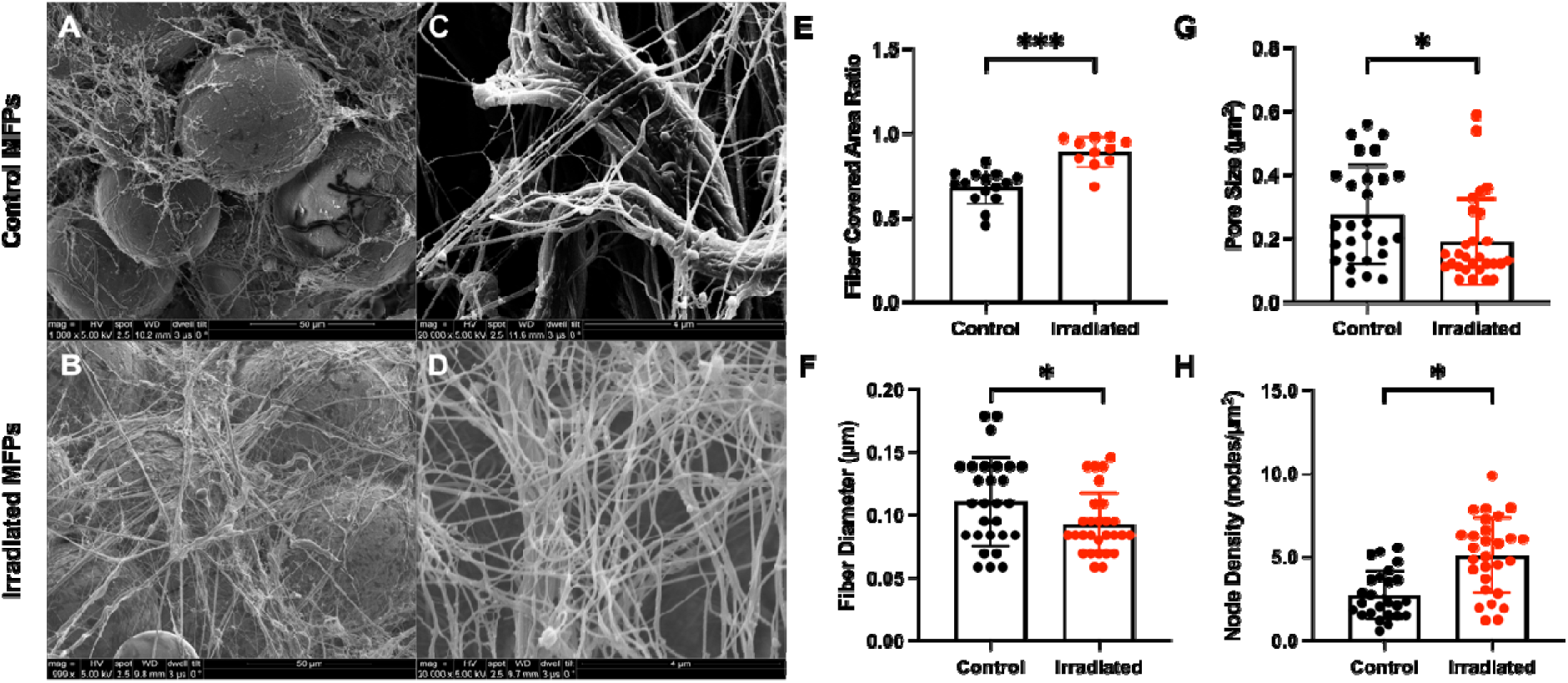
Structure of control and irradiated MFPs by SEM and fiber network analysis. SEM images of unirradiated control (0 Gy) (**A**) and *ex vivo* irradiated (20 Gy) (**B**) MFPs from Nu/Nu mice (n = 10 mice) at 1,000x, and control (**C**) and irradiated (**D**) MFPs at 20,000x magnification. SEM images were analyzed using an automated fiber tracking algorithm to determine the fiber covered area ratio (**E**), average fiber diameter (**F**), pore size (**G**), and node density (**H**). Statistical significance was determined by Student’s t-test with *p<0.05 and ***p<0.001. Error bars show standard deviation.

We then used Raman spectroscopy to evaluate the biochemical composition changes following MFP irradiation (**Supplementary Fig. S2A**). The peaks at 1300 cm^-1^ are from lipids, which are responsible for vibrations of -(CH_2_)_n_-in-plane twist and fatty acids [23,24]. The peak at 1265 cm^-1^ is generally assigned to proteins (Amide III), and collagen is a major contributor of the vibrational motion. The 1650 cm^-1^ area is a broad peak with multiple contributions, including proteins and Amide I [25,26]. Taken together, these peaks have a strong likelihood of being contributed by collagen in tissues [25–30]. **Supplementary Figs. S2B-C** show that collagen/lipid ratios (both amide I/lipid and amide III/lipid) increased significantly in MFPs (p<0.01) after RT, confirming the increase in ECM deposition observed in SEM images. In an effort to elucidate the biochemical changes between irradiated and control samples, we used PCA to simplify the complexity of high-dimensional data while retaining relevant information necessary for classification [31–33]. In a two-dimensional PC scatter plot, the first principal component (PC1) and the second principal component (PC2) showed a clear separation between the control and irradiated groups with a variance level of 29.36% for PC1 and 9.66% for PC2, again confirming compositional changes induced by RT (**Supplementary Fig. S2D**). Normalized Raman spectra peak analysis validated this difference, especially for the collagen peaks (**Supplementary Table S2**).

### 3.2 Mechanical properties and ECM component analysis

SEM images and Raman spectroscopy results indicated that ECM deposition was enhanced in irradiated MFPs. To characterize the impact of increased ECM fibers on tissue biomechanical properties, AFM was used to determine the stiffness of MFPs. FD measurements (**Supplementary Fig. S3**) were taken to determine Young’s moduli [34]. The range of elastic moduli for control and irradiated breast tissue is shown in **Fig. 2A** and **Fig. 2B**. The histogram of stiffness values from control MFPs reveals a bimodal stiffness distribution with two prominent peaks at 0.2 kPa (‘peak 1’) and 0.6 kPa (‘peak 2’) (**Fig. 2A**). In comparison, irradiated MFPs exhibit a bimodal stiffness distribution with two prominent peaks at 0.4 kPa (‘peak 1’) and 1 kPa (‘peak 2’). At values higher than 2 kPa, a broadening in the distribution indicates a marked mechanical heterogeneity across the sample. The average Young’s modulus for irradiated MFPs was 2.4-fold greater than control MFPs (**Fig. 2B**).

**Figure 2.**
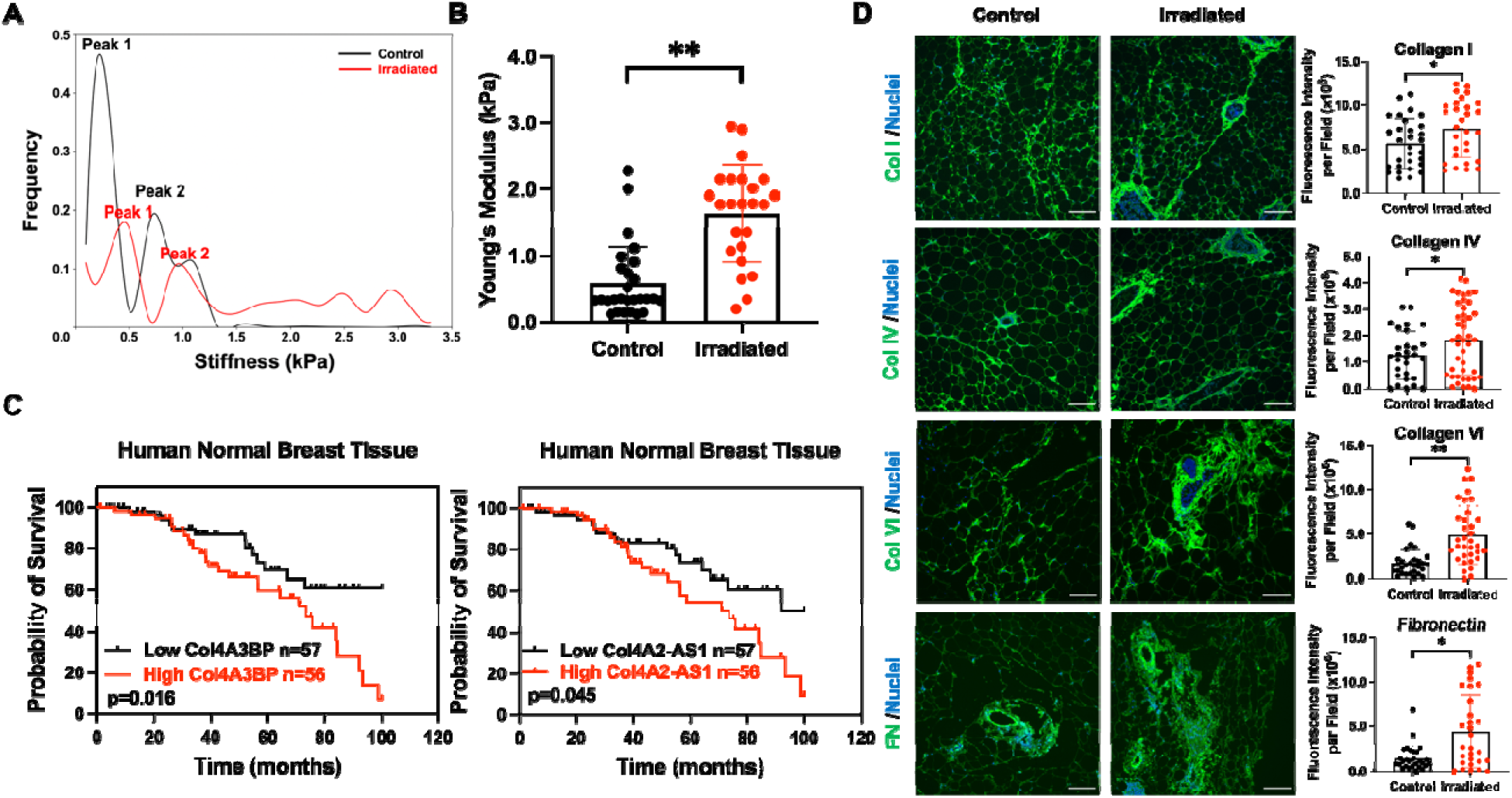
MFP irradiation enhances tissue stiffness and ECM deposition. **(A)** Stiffness distribution of control and *ex vivo* irradiated (20 Gy) MFPs from Nu/Nu mice (n = 4 mice). (**B**) AFM analysis of the Young’s modulus of control and *ex vivo* irradiated MFPs using the Hertz model. (**C**) Kaplan-Meier curves for patients with low versus high mRNA expression of *Col4A3BP* and *Col4A2-AS1* (n = 113 patients) in the breast tumor-adjacent normal tissue from the TCGA database. Immunofluorescence (IF) was performed to detect collagen I (Col I), collagen IV (Col IV), collagen VI (Col VI), and fibronectin (FN) in MFPs, and representative images and the corresponding quantification of fluorescence intensity is shown in (**D**). Scale bars are 100 μm. Statistical significance was determined by Student’s t-test with *p<0.05 and **p<0.01. Error bars show standard deviation.

To determine how ECM components in normal breast tissue impact patient outcomes, we obtained mRNA expression data from the normal adjacent tissue of patients with breast cancer from TCGA datasets. We found that high mRNA expression of collagen type IV alpha-3-binding protein *(Col4A3BP)* and collagen type IV alpha-2 antisense RNA 1*(Col4A2-AS1)* were correlated with poor overall survival (**Fig. 2C**). *Col4A3BP* phosphorylates the non-collagenous domain of the α3 chain of collagen IV and is associated with epithelial-to-mesenchymal transition and mesh-like collagen IV formation [35] while *Col4A2-AS1* is a long non-coding RNA for collagen IV α2 that is associated with the growth and metastasis of breast cancer cells [36].

These associations led us to investigate specific changes in ECM components in irradiated breast tissue. IF staining showed that collagen I, collagen IV, collagen VI, and fibronectin (FN) expression were increased in irradiated murine breast tissue (**Fig. 2D**). Masson’s trichrome staining confirmed an increase in overall collagen in irradiated MFPs (**Supplementary Fig. S4**). These data illustrate that RT modulates the expression of ECM subcomponents and leads to changes in ECM deposition and stiffness, which may influence tumor behavior and negatively impact patient survival in breast cancer.

### 3.3 dECM hydrogels

*MFP decellularization:* Motivated by the RT-induced increase in ECM deposition that may provide a platform for cell adhesion and invasion, we next utilized a decellularization technique to extract ECM, develop dECM hydrogels, and encapsulate cells to observe their behavior. We used an immunodeficient model to replicate *in vivo* conditions relevant to recurrence [6]. Murine MFPs were decellularized by a series of mechanical, chemical, and enzymatic methods [15]. The final volume of decellularized MFPs was 10–20% of the original adipose tissue. H&E staining confirmed the removal of cell nuclei while preserving ECM proteins following decellularization (**Fig. 3A, Supplementary Fig. S5**). We also verified the retention of ECM using Masson’s trichrome staining as well as lipid removal using perilipin staining. IF staining showed that decellularization of MFPs retained post-RT trends in collagen I, collagen IV, collagen VI, and FN expression (**Supplementary Fig. S6**). The above results indicate that the extracellular components were preserved while the cells and lipids were removed from decellularized MFPs.

**Figure 3.**
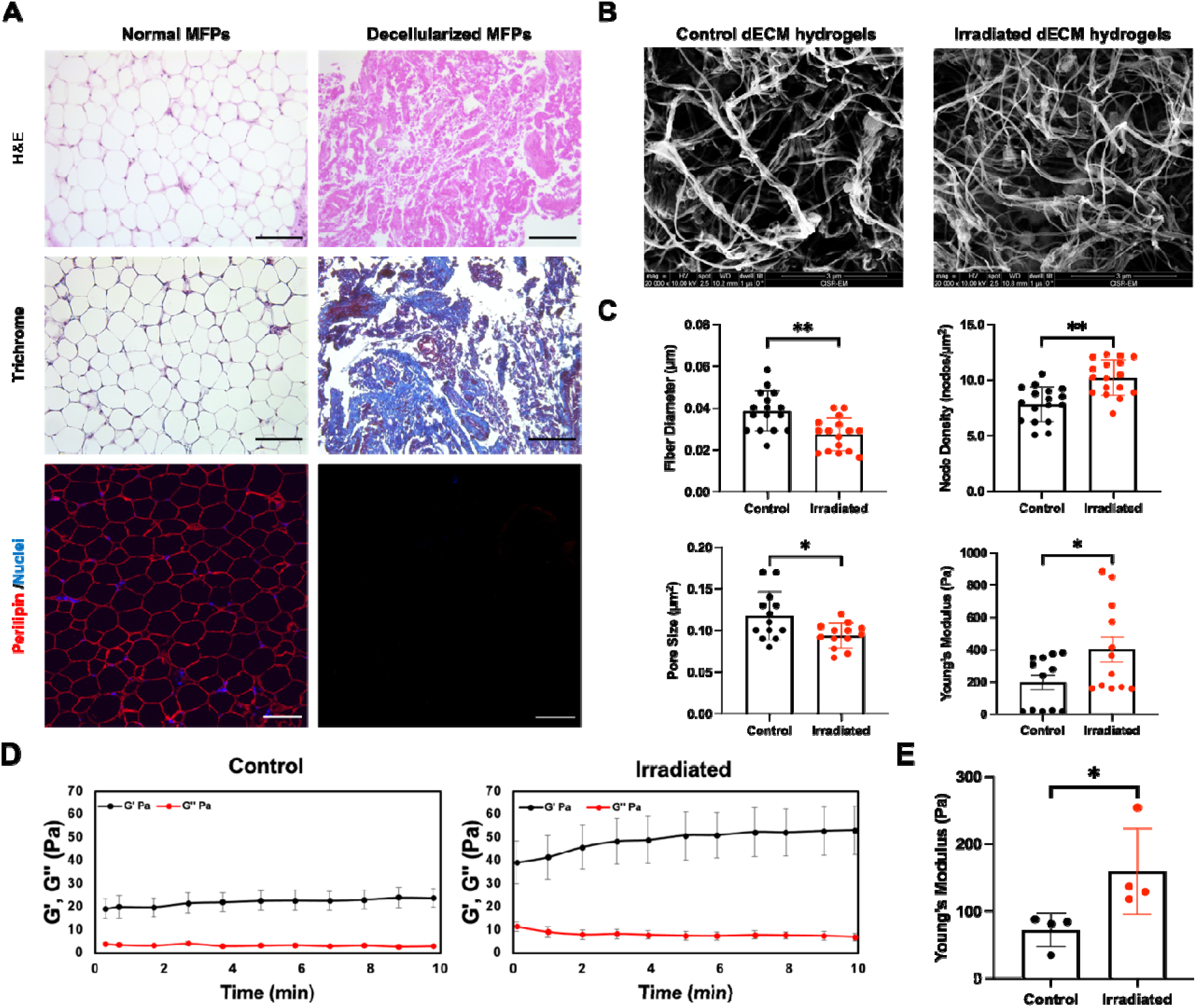
Characterization of dECM hydrogels from Nu/Nu mice. (**A**) H&E, Masson’s trichrome, perilipin (red), and nuclear (blue) staining of pre- and post-decellularized control MFPs confirmed decellularization and delipidation. Scale bars are 100 μm. SEM images of control and irradiated dECM hydrogels at 20,000x magnification (**B**), and the average fiber diameter, node density, pore size, and mechanical properties determined by AFM were evaluated (**C**). (**D**) Time sweep rheology analysis is shown for the storage (G’) and loss modulus (G’’) of control and irradiated dECM hydrogels. (**E**) Young’s moduli were quantified for control and irradiated dECM hydrogels from rheology analysis. Statistical significance was determined by Student’s t-test with *p<0.05 and **p<0.01 (n = 4 biological replicates). Error bars show standard deviation.

*dECM hydrogel characterization:* To evaluate the structural and stiffness characteristics of control and irradiated dECM hydrogels, SEM and AFM were performed. SEM was utilized to determine the architecture of dECM hydrogels without cell encapsulation (**Fig. 3B**). Automatic fiber quantification of SEM images showed the parameters of fiber diameter, node density, and pore size were altered in irradiated dECM hydrogels (**Fig. 3C**), which parallels the changes observed in irradiated MFPs. These differences are likely due to the varying collagen concentration post-RT [37,38]. Furthermore, AFM measurements showed that irradiated dECM hydrogel stiffness was 2-fold higher than those for control dECM hydrogels. We also employed rheology to determine the mechanical properties of dECM hydrogels. The storage modulus (G’) and loss modulus (G’’) of dECM hydrogels derived from RT-treated mice exhibited a notable increase (**Fig. 3D**), and a 2-fold increase in the Young’s modulus of irradiated dECM hydrogels was again observed (**Fig. 3E**). Together, these data demonstrate the integrity of ECM after decellularization and that the changes in dECM hydrogels between control and irradiated samples are consistent in control and irradiated MFPs.

### 3.4 Proliferation and invasion of breast cancer cells in dECM hydrogels

We encapsulated 4T1 murine TNBC cells in control and irradiated dECM hydrogels and evaluated their response after 48 hr (**Fig. 4A**). A Live/Dead viability assay showed that dECM hydrogels are not cytotoxic to 4T1 cells (**Supplementary Fig. S7A-C**). Additionally, IF and bioluminescence imaging were used to quantify the proliferation of GFP- and luciferase-labeled 4T1 cells in dECM hydrogels (**Fig. 4B**). 4T1 cell proliferation increased significantly in irradiated dECM hydrogels compared to unirradiated controls (p<0.05). MDA-MB-231 human TNBC cells also showed enhanced proliferation in the irradiated microenvironment (**Supplementary Fig. S8A-C**), suggesting that irradiated dECM hydrogels promote tumor cell proliferation. To investigate the impact of the irradiated microenvironment on 4T1 invasiveness, we conducted a modified transwell invasion assay. We found that 4T1 cells encapsulated in irradiated dECM hydrogels displayed significantly increased invasion when compared with cells interacting with control dECM hydrogels (**Fig. 4C**).

**Figure 4.**
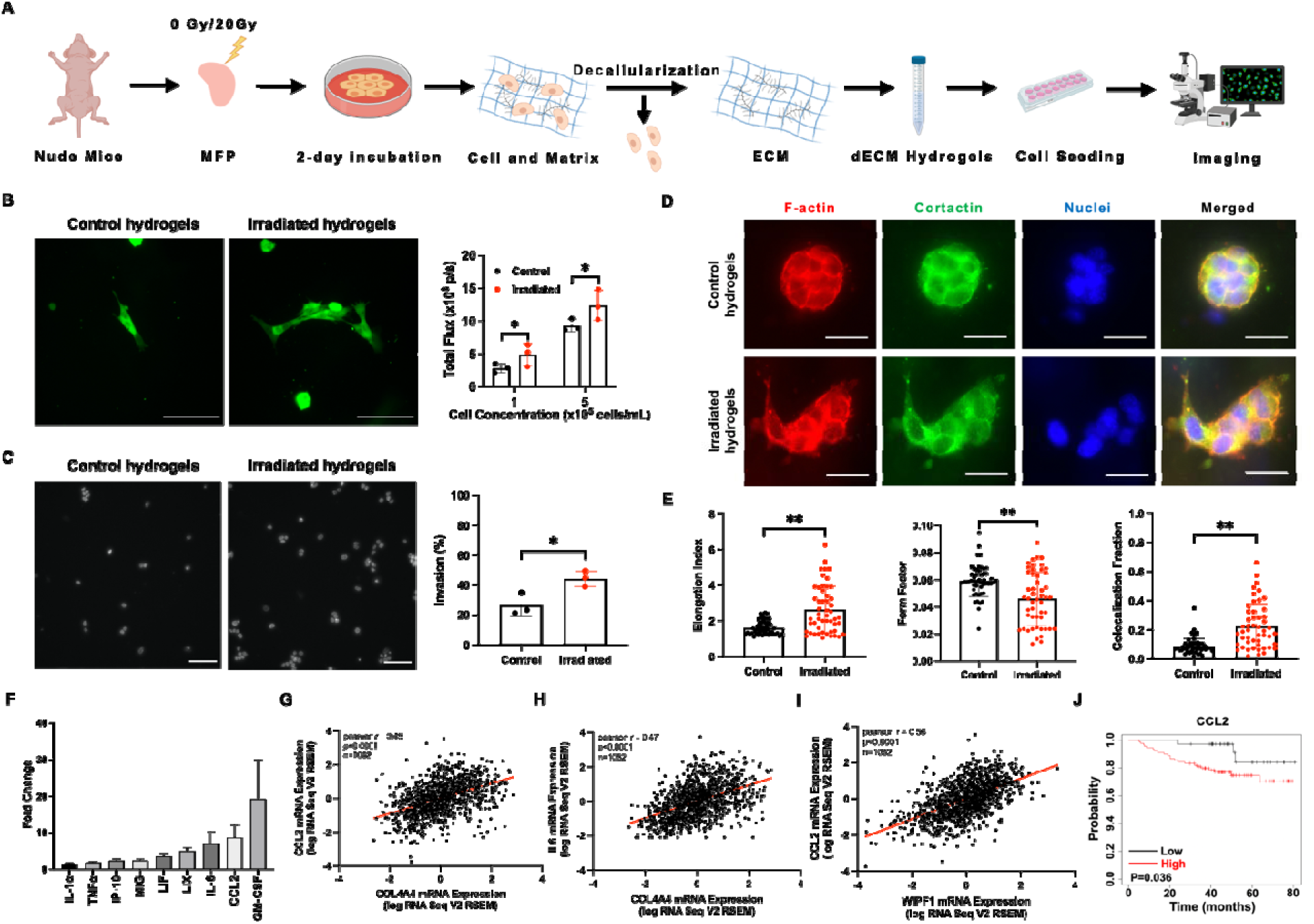
The irradiated microenvironment promotes tumor cell proliferation and invasion and worse outcomes in TNBC patients. (**A**) Schematic of experimental workflow. (**B**) Fluorescence images of GFP- and luciferase-labeled 4T1 cells in dECM hydrogels from Nu/Nu mice (n = 3 for each condition) and bioluminescence imaging quantification. Scale bars are 100 μm. (**C**) Representative nuclei images of invading 4T1 cells and quantification in control and irradiated dECM hydrogels. Scale bars are 100 μm. (**D**) Staining of F-actin (red), cortactin (green), nuclei (blue) and merged images for 4T1 cells embedded in control and *ex vivo* irradiated dECM hydrogels for 48 hr. Scale bars are 50 μm (**E**) Cellular elongation index and form factor based on F-actin staining and cortactin/F-actin colocalization as determined by fractional pixel overlap were analyzed (n = 3 biological replicates). (**F**) A Luminex multiplex immunoassay was performed to determine the secreted cytokine profile of CM from 4T1 cells cultured in irradiated dECM hydrogels (n = 3). Data presented is mean fluorescence intensity (MFI) fold change compared to cytokine secretion from 4T1 cells cultured in control dECM hydrogels. Only factors with a 1.5-fold or greater increase in MFI are shown. **Supplementary Table S1** defines abbreviations. Correlation of *Col4A4* and *CCL2* mRNA expression (**G**), *Col4A4* and *IL6* mRNA expression (**H**), and *WIPF1* and *IL6* mRNA expression (**I**) from TCGA analysis (PanCancer dataset) in 1,082 patients with breast tumors. Pearson correlation shown. (**J**) Kaplan-Meier curves of TNBC patients (n = 126 patients) with low versus high mRNA expression of *CCL2* from KMplotter. Statistical significance was determined by Student’s t-test with* p<0.05, **p<0.01 and ***p<0.001. Error bars show standard deviation.

To verify the upregulated invasion of TNBC cells, we further visualized the cytoskeletal properties of 4T1 cells embedded in the dECM hydrogels. The driving force for cancer invasion into their surrounding microenvironment as well as numerous cellular physical processes is through localized actin filaments [39–42]. Therefore, we stained for F-actin using phalloidin, and cellular morphology was quantified in the irradiated dECM hydrogels (**Fig. 4D-E**). Elongation index and form factor can be used as measures of cell morphology, where high elongation index and low form factor indicate higher invasiveness [43,44]. Cells in the irradiated group increased their elongation index (p<0.05) and reduced their form factor significantly (p<0.05), suggesting cell invasiveness increases in the irradiated microenvironment. We also examined invadopodia, which are protrusive structures that can remodel the ECM and facilitate invasive migration [45–47]. The colocalization of F-actin and cortactin are considered reliable indicators for invadopodia [48]. F-actin and cortactin colocalization revealed significantly more invadopodia in 4T1 cells encapsulated within irradiated hydrogels (p<0.05), indicating a higher invasive capacity of 4T1 cells influenced by an irradiated microenvironment (**Fig. 4D-E**) and confirming the transwell assay result. Tyrosine kinase substrate 4 (TKS4) and WASP interacting protein (WIP) are key molecular components that modulate invadopodia extension [49]. Clinical data from TCGA showed an association between the gene expression of collagen subtypes IV and VI and TKS4 (*SH3PXD2B*) or WIP (*WIPF*) in patients with breast cancer (**Supplementary Fig. S9A-B**), indicating high invadopodia formation may be the result of increased microenvironmental collagen expression.

We next collected CM from 4T1 cells encapsulated in the dECM hydrogels and analyzed cytokine secretion using a Luminex multiplex immunoassay. Factors with a 1.5-fold or greater increase in mean fluorescence intensity (MFI) are shown (**Fig. 4F**), with granulocyte macrophage colony-stimulating factor (GM-CSF), CCL2, and interleukin 6 (IL-6) having the highest expression. These cytokines are associated with an increase in tumor cell invasiveness [50,51]. To understand the relationship between ECM components, invadopodia, and cytokine secretion in a broader clinical context, TCGA analysis revealed a correlation between *Col4A4* and *CCL2* mRNA expression (**Fig. 4G**). We also found a correlation between *Col4A4* and *IL6* expression (**Fig. 4H**), indicating the increased collagen IV levels in breast tissue after RT may elevate the secretion of cytokines in tumor cells. Further analysis of the TCGA PanCancer dataset showed *WIPF1* was associated with *CCL2* mRNA expression (**Fig. 4I**). This clinical data suggests a link between increased collagen IV deposition in the tumor microenvironment, invadopodia regulation, and CCL2 expression, which mirrors our findings that tumor cell cytokine secretion is enhanced in response to the increase in collagen IV after RT. To investigate the link between CCL2 secretion and patient outcomes in TNBC, we utilized publicly available datasets through KMplotter. Among TNBC patients, high CCL2 expression was associated with poor patient survival (**Fig. 4J**). These data indicate that RT-induced collagen IV deposition may lead to elevated levels of CCL2 in recruited tumor cells, ultimately decreasing patient survival.

### 3.5 Enhanced tumor proliferation and M2 polarization by 4T1 cell and BMDM interactions in dECM hydrogels

Both GM-CSF and CCL2 expression are associated with macrophage infiltration [52], and GM-CSF has specifically been implicated in post-RT macrophage and circulating tumor cell infiltration [5,6]. Macrophages are notably abundant in tumors and fulfill crucial roles at various stages of tumor progression [53]. We therefore evaluated how macrophages behave in irradiated dECM hydrogels and how they interact with tumor cells. We isolated BMDMs from Nu/Nu mouse femurs and seeded them in dECM hydrogels. Ki67 staining showed that BMDMs increased their proliferation in the irradiated microenvironment (**Fig. 5A-B**). This increased BMDM proliferation was also confirmed using an alamarBlue assay (**Supplementary Fig. S10A-B**). We then co-cultured GFP-labeled 4T1 cells and BMDMs stained with CellTrace Far Red to evaluate differences in their interactions within irradiated and unirradiated microenvironments. Fluorescence images and bioluminescence measurements demonstrated an increase in 4T1 cell proliferation when co-cultured with macrophages in the irradiated hydrogels (**Supplementary Fig. S10C**), which was confirmed by Ki67 staining (**Fig. 5C-E**). Notably, this increased proliferation is 2-fold higher when compared to the proliferation of 4T1 cells cultured alone (**Fig. 4B**), suggesting that macrophages further promote tumor cell proliferation in irradiated microenvironments. BMDMs also exhibited increased proliferation in the irradiated microenvironment when co-cultured with 4T1 cells (**Fig. 5C-E**).

**Figure 5.**
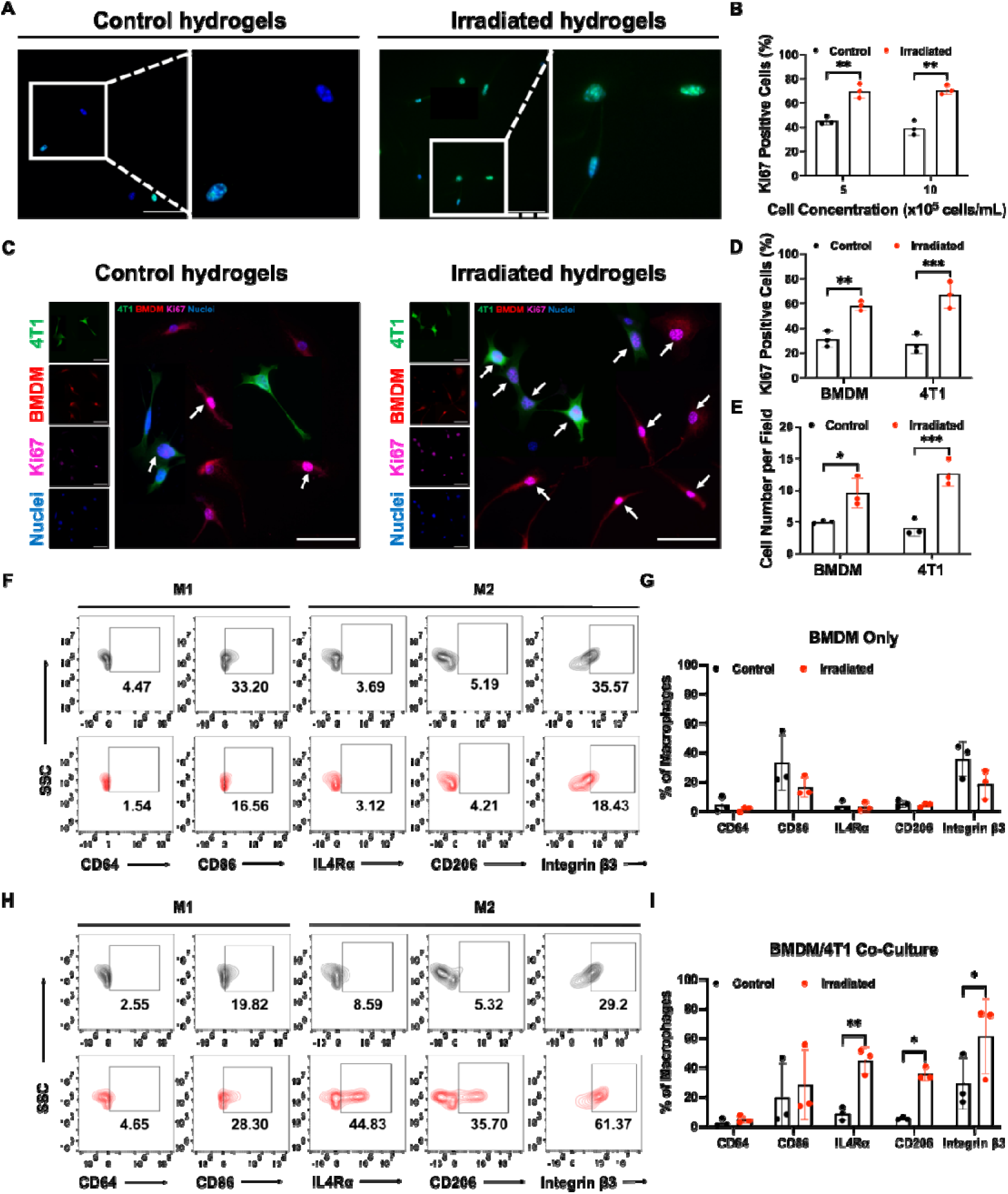
Tumor-macrophage interactions promote tumor cell proliferation and M2 macrophage polarization in irradiated dECM hydrogels. (**A**) Fluorescence images of Ki67 positive (green) BMDMs in control and irradiated dECM hydrogels with the corresponding quantification in (**B**). Nuclei are stained blue. (**C**) Representative images of Ki67 staining (purple) in co-cultures of GFP-labeled 4T1 cells (green) and BMDMs (red) stained with CellTrace Far Red in control and irradiated dECM hydrogels. White arrows indicate positive Ki67 staining. The percentage of Ki67 positive cells (**D**) and cell number per field (**E**) for 4T1 cells and BMDMs were quantified. Flow cytometry analysis (**F**) and quantification (**G**) of M1 markers (CD64/CD86), M2 markers (IL4R/CD206), and integrin β3 of F4/80^+^ BMDMs cultured alone in control and irradiated dECM hydrogels. Flow cytometry characterization (**H**) and quantification (**I**) of F4/80^+^ BMDMs co-cultured with 4T1 cells in control and irradiated dECM hydrogels. Scale bars are 50 μm. Statistical significance was determined by Student’s t-test with *p<0.05, **p<0.01, and ***p<0.01. Error bars show the mean with standard deviation.

Macrophages have the capacity to adopt distinct functional phenotypes, known as M1 and M2 macrophages [54]. M1-like macrophages are characterized as pro-inflammatory cells responsible for eliminating cancer cells, whereas M2-like macrophages are anti-inflammatory cells that foster tumor growth, promote metastasis, facilitate tissue remodeling, and induce immunosuppression [54–56]. We sought to investigate macrophage phenotype shifts in dECM hydrogels. We characterized the phenotypes of F4/80^+^ macrophages as follows: macrophages exhibiting elevated levels of CD86 and CD64 were classified as pro-inflammatory M1 macrophages while those displaying increased expression of IL4Rα and CD206 were categorized as anti-inflammatory M2 macrophages (**Supplementary Fig. S11A-C**). Integrin β3 is a cell surface receptor protein that plays an important role in cell-cell adhesion, cell-ECM interactions, and cell signaling [57]. The recruitment of macrophages is partially regulated by integrin β3, and the expression of integrin β3 on the surface of macrophages is associated M2 polarization [58–60]. Flow cytometry analysis revealed that there were no statistically significant differences in the expression levels of M1 and M2 markers among BMDMs cultured alone in both control and irradiated dECM hydrogels (**Fig. 5G**). However, the BMDM phenotype shifted significantly toward an M2 phenotype with upregulated integrin β3 expression when co-cultured with 4T1 cells in the irradiated dECM hydrogels (**Fig. 5H, Supplementary Fig. S11D**). We propose that the increase in stiffness in the irradiated microenvironment directly influences tumor-macrophage interactions to promote M2 macrophage polarization and tumor cell proliferation and invasion (**Fig. 6**).

**Figure 6.**
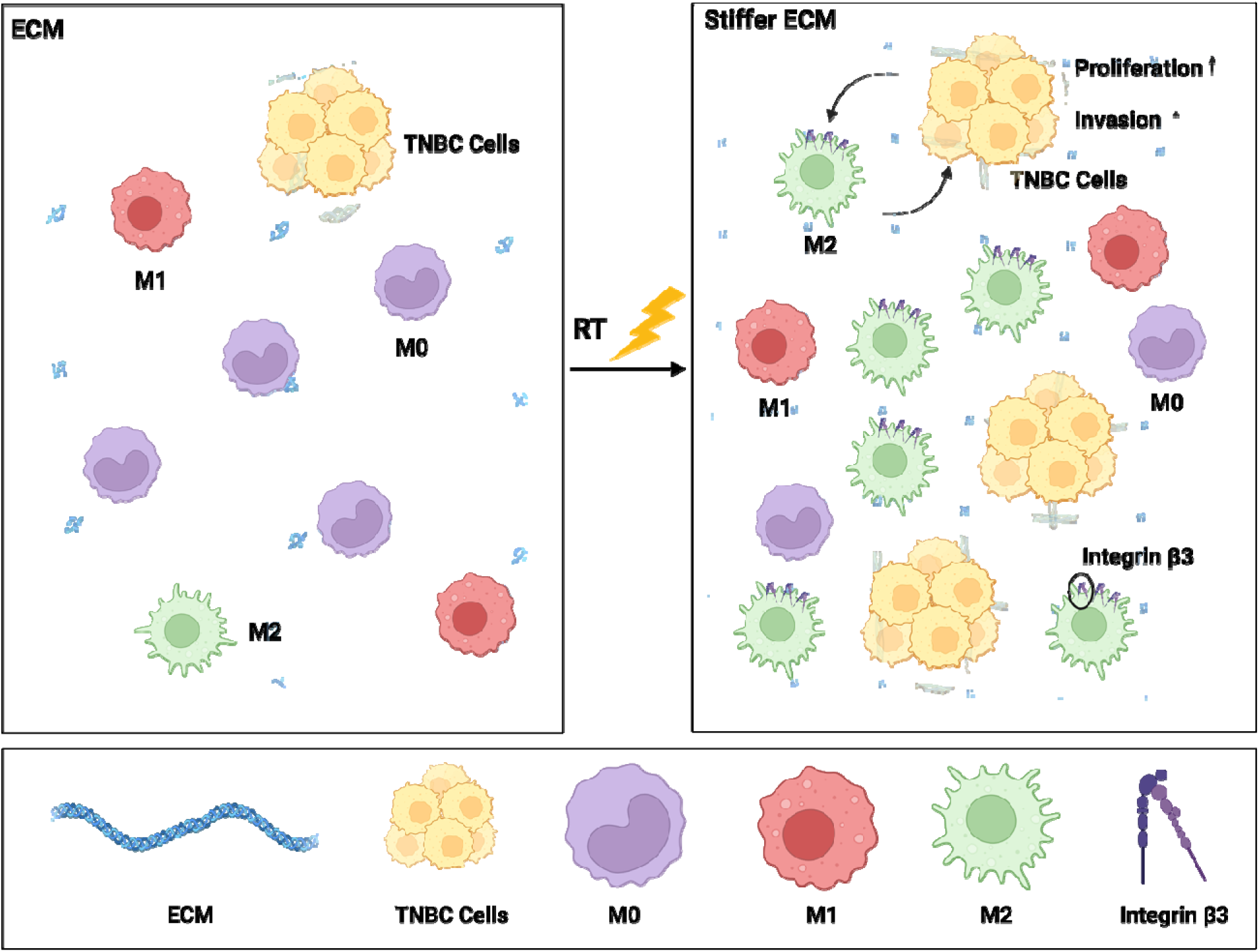
Model of pro-tumor ECM changes driven by radiation in the breast microenvironment. Radiation-induced ECM changes promote tumor-macrophage interactions that enhance tumor cell invasion and proliferation linked to immunosuppressive M2 macrophage polarization.

## 4. Discussion

The high incidence of local recurrence following RT is associated with poor overall survival in TNBC patients, and previous work suggests a link between radiation damage and TNBC recurrence in immunocompromised patients and mouse models [6,61]. Understanding how irradiation modulates the microenvironment is necessary for evaluating drivers of recurrence. We first analyzed structural and compositional alterations of irradiated breast tissue and showed more ECM deposition, reduced fiber diameter and pore size, and increased stiffness following irradiation. We determined that collagen IV-associated changes in normal tissues predicted worse outcomes in breast cancer patients, and we confirmed an increase in multiple ECM components, including collagen IV, in MFPs post-RT. As physicochemical cues dictated by changes in ECM deposition modulate numerous biological functions, we developed dECM hydrogels as models of the irradiated microenvironment to determine the impact of RT-induced ECM changes on tumor cell behavior. We showed that normal tissue radiation damage leads to an environment favorable for TNBC recurrence and M2 phenotype polarization. Our study is the first to recapitulate the irradiated ECM microenvironment, and this model can be used as a platform for studying radiation response and recurrence mechanisms.

The recruitment and transformation of fibroblasts results in dysregulated ECM remodeling after immune cell infiltration to injured sites, and prolonged ECM remodeling can contribute to the development of fibrosis [9,62–64]. The ECM in injured tissue is usually denser and mechanically stiffer than normal ECM, and the physical characteristics of the newly deposited fibers are critical in affecting how cells sense their microenvironment [65–67]. These findings are consistent with our SEM results showing excess ECM deposition and denser and thinner ECM fibers in irradiated samples, and our AFM and rheology studies confirmed an increase in tissue stiffness due to the altered ECM post-RT. The stiffness range of human breast tissue is approximately 0.5 - 1 kPa [68], which is in the range of our findings. RT may activate fibroblasts, recognized as stromal regulators, which in turn play a pivotal role in ECM modification [9,69–71]. Notably, the thinner fibers and enhanced mechanical properties observed could be linked to increased aberrant ECM deposition post-RT [72–74]. Recent studies also showed that increased ECM cross-linking induced by high levels of lysyl oxidases may elevate tissue stiffness [69,75,76]. Accordingly, increased tissue stiffness is influenced by ECM component synthesis, assembly, and cross-linking in the irradiated microenvironment, which has the potential to impact tumor progression.

Changes in ECM density and stiffness are not just attributed to fiber size but are also impacted by the molecular makeup of the matrix [66]. Approximately 300 proteins present in the ECM have been reported to mediate cellular behaviors and tissue development, and collagen is the major fibrous protein which constitutes up to 90% of the ECM [77]. 28 types of collagen have been described [78], and collagens I, IV, and VI are the most relevant subtypes of collagens in the microenvironment of mammary tissue. A major function of collagens I, IV, and VI is to provide structural support for cells and enrich the cell surface with growth factors and cytokines that are important in tumor progression. Increased collagen IV after chemotherapy has been shown to drive TNBC cell invasion [79]. Collagen VI is secreted by adipocytes and is known to regulate breast tumor development and facilitate TNBC cell migration [80,81]. In addition to collagens, other prominent fibrous proteins including elastins, fibronectins, and laminins are related to cellular dynamics [82]. Fibronectins are crucial for attaching cells to matrix and guiding cell migration and have been reported to be upregulated in the RT-induced fibrosis process [83]. Radiation damage is known to alter the expression of ECM components in addition to ECM modifying enzymes [84–86], which agrees with our results showing an increase in ECM component expression and remodeling in irradiated MFPs. We also showed that changes in collagen deposition may provide more tensile strength and support tumor cell retention and invasion through direct structural changes and cell-ECM contact.

Variations in ECM density, composition, and stiffness have been correlated with tumor progression and recurrence [82]. In this study, we aimed to isolate the specific impact of post-RT changes in ECM on tumor and immune cell behavior while replicating *in vivo* conditions through the development of irradiated mammary tissue-derived dECM hydrogels. Collagen- and Matrigel-based hydrogels have been widely used to study cell behavior, but they have limitations due to the absence of native tissue structure or site-specific bioactive molecules such as exogenous growth factors and peptides [87,88]. The advantages of component integrity and biologically active constituent matrix molecules make dECM hydrogels a promising method to mimic the irradiated microenvironment [89]. Our results confirm the preservation of ECM components following the decellularization process to form hydrogels. dECM hydrogels have been previously fabricated from various tissue sources, including adipose tissue [13,90], liver [91], lung [92], dermis, and urinary bladder [12], with hydrogel mechanical properties ranging from 0.1 to 40 kPa [12,93], consistent with our results. The pore size, fiber density, and fiber orientation in dECM hydrogels have previously been demonstrated to lead to variations in mechanical properties [94,95]. Dermal dECM hydrogels have a higher fiber node density than urinary bladder dECM hydrogels and a higher storage modulus, indicating a link between fiber node density and mechanical strength [96]. Our SEM images and fiber analysis showed that fibers in irradiated dECM hydrogels were denser compared to unirradiated dECM hydrogels, which correlated to increased stiffness. Changes in ECM stiffness are known to impact cell motility [97], and we observed that the increased stiffness in dECM hydrogels after radiation had a profound effect on tumor cell invadopodia formation and cellular morphology indicative of increased invasiveness. Tissue-specific ECM has also been shown to influence tumor cell growth [98] directly through engagement with integrins [99], and we found that tumor cell proliferation increased when encapsulated within irradiated dECM hydrogels. We determined that breast cancer cell interactions with the irradiated ECM promote a pro-tumor microenvironment through increased proliferation and invasion, which builds on previous findings linking the recruitment of tumor cells to irradiated sites to recurrence under immunocompromised conditions [6].

Immune cells are recruited to RT-damaged sites by inflammatory mediators and secrete profibrotic cytokines such as tumor necrosis factor alpha, IL-13, and transforming growth factor-β during fibrosis [9]. Macrophages are known to infiltrate irradiated normal tissue [100], and the presence of inflammatory macrophages has been correlated to recurrence and poor overall survival in breast cancer patients [101]. Tumor-associated macrophages (TAMs) initially exhibit an M1-like phenotype during the early stages of tumor progression while predominantly shifting toward to M2-like phenotype during tumor growth [54–56]. The M2 macrophage phenotype is well recognized to support cancer metastasis by mediating genes involved in immunosuppression, cytoskeletal remodeling, and angiogenesis [54,102]. For example, M2a and M2c macrophages promote lung cancer proliferation and invasion. However, M1 macrophages suppress lung tumor growth [103]. Additionally, clinical data has shown higher infiltration of TAMs, especially with an M2 phenotype, is associated with worse overall patient survival in multiple cancer types, including breast, lung, and bladder cancer [103–106]. Previous literature also indicates that infiltrating macrophages polarize to an M2 phenotype in irradiated breast tissue and contribute to tumor recurrence [6,7]. We did not model macrophage recruitment in this study but instead showed that the irradiated microenvironment supports macrophage growth. Interestingly, we found that macrophages shifted toward an M2 phenotype in irradiated dECM hydrogels when they interacted with 4T1 cells, and this interaction also increased 4T1 proliferation. Although increases in tumor cell proliferation occurred without macrophages in irradiated dECM hydrogels, the extent of cancer cell proliferation was greater during co-culture with macrophages. This suggests that the irradiated ECM influences tumor-immune cell interactions and stimulates the macrophage phenotype transition to create an immunosuppressive, pro-tumor, and pro-recurrent niche. This agrees with previous *in vivo* findings showing that excess macrophage infiltration attracts circulating tumor cells and indicates that cross-talk with immune cells plays a pivotal role in tumor cell survival in irradiated tissues [5,6,107].

In our study, we also found that high expression of M2 markers was accompanied by elevated levels of integrin β3, which mediates cell-cell adhesion, cell-ECM interactions, and intracellular signaling pathways [58,108]. The abnormal expression of numerous integrins is reported to regulate cell-ECM interactions and facilitate malignant progression in breast cancer patients [58,109]. Additionally, recent studies show that stiff ECM cooperates with integrin β3 to promote ErbB2-dependent breast cancer progression by elevated IR/AKT/mTORC1 signaling, suggesting altered integrin β3 in response to tissue stiffness [110]. Previous work has also shown that M2 polarization is dependent on integrin β3 through the PPARγ pathway [59]. Together, we propose a potential underlying mechanism that tumor-macrophage interactions stimulate the activation of integrin β3 and enhance M2 polarization in stiffer ECM induced by RT, and M2 macrophages subsequently increase tumor cell proliferation.

Overall, we developed a mammary-derived dECM hydrogel to model the normal tissue radiation response, which will lead to the discovery of novel targets for reducing TNBC recurrence and increasing breast cancer patient survival following therapy. We replicated *in vivo* conditions using dECM hydrogels to study the influence of the irradiated breast tissue on TNBC and immune cell behavior. Although our MFPs were irradiated *ex vivo*, which may have different outcomes compared to *in vivo* irradiation where cell infiltration may further alter ECM remodeling, this model is necessary for analyzing the direct effect of radiation on changes in the ECM within mammary tissue. In addition, post-RT breast biopsies are not part of the standard of care for TNBC patients, and thus *ex vivo* irradiation provides a valuable point of comparison with any patient samples that may be studied. Future studies will evaluate the impact of *in vivo* RT in mice on ECM-linked TNBC recurrence.

## 5. Conclusions

We demonstrated that dECM hydrogels are an effective tool to study the normal tissue radiation response. ECM structural and compositional changes induced by radiation facilitated a pro-tumor niche through increased tumor cell proliferation and invasion. In addition, we showed that the irradiated microenvironment enhanced immunosuppressive M2 macrophage polarization that further stimulated TNBC cell proliferation, which may drive TNBC recurrence following therapy.

## Acknowledgments

The authors thank Dr. Laura L. Bronsart for providing the GFP- and luciferase-labeled 4T1 cells, Dr. Michael Freeman for irradiator use, Dr. Craig L. Duvall for IVIS and lyophilizer use, and Dr. Scott Guelcher for rheometer use. We also thank the Vanderbilt Cell Imaging Shared Resource (CISR) core for SEM imaging, the Translational Pathology Shared Resource (TPSR) for Raman sample and IF sample preparation, and Dr. Dmitry S. Koktysh in the Vanderbilt Institute of Nanoscale Science and Engineering (VINSE) core for assistance with AFM measurements. The results shown here are in part based upon data generated by the TCGA Research Network: https://www.cancer.gov/tcga. We appreciate Dr. Runchen Zhao’s F-actin-cortactin co-localization MATLAB code and Co Quach Dai’s stiffness frequency distribution figure preparation. This research was financially supported by NIH grants #R00CA201304, F31CA254311 (BCH), S10RR026373 (CSIR), and P30CA068485 (TPSR) as well as the Concern Foundation Conquer Cancer Now Award, the American Cancer Society Research Scholar Grant, and a VINSE pilot award. Schematics were created with BioRender.com.

## Data Availability

Data will be made available upon request.

## References

[1] Breast cancer facts & figures 2022–2024, Am. Cancer Soc. (2022) 1–48.

[2] N.L. Simone, T. Dan, J. Shih, S.L. Smith, L. Sciuto, E. Lita, M.E. Lippman, E. Glatstein, S.M. Swain, D.N. Danforth, K. Camphausen, Twenty-five year results of the national cancer institute randomized breast conservation trial, Breast Cancer Res. Treat. 132 (2012) 197–203. 10.1007/s10549-011-1867-6.

[3] B.G. Haffty, Long-term results of hypofractionated radiation therapy for breast cancer, Breast Dis. 21 (2010) 267–268. 10.1016/S1043-321X(10)79594-1.

[4] A.J. Lowery, M.R. Kell, R.W. Glynn, M.J. Kerin, K.J. Sweeney, Locoregional recurrence after breast cancer surgery: a systematic review by receptor phenotype, Breast Cancer Res. Treat. 133 (2012) 831–841. 10.1007/s10549-011-1891-6.

[5] M. Vilalta, M. Rafat, A.J. Giaccia, E.E. Graves, Recruitment of circulating breast cancer cells is stimulated by radiotherapy, Cell Rep. 8 (2014) 402–409. 10.1016/j.celrep.2014.06.011.

[6] M. Rafat, T.A. Aguilera, M. Vilalta, L.L. Bronsart, L.A. Soto, R. Von Eyben, M.A. Golla, Y. Ahrari, S. Melemenidis, A. Afghahi, M.J. Jenkins, A.W. Kurian, K.C. Horst, A.J. Giaccia, E.E. Graves, Macrophages promote circulating tumor cell-mediated local recurrence following radiotherapy in immunosuppressed patients, Cancer Res. (2018). 10.1158/0008-5472.CAN-17-3623.

[7] B.C. Hacker, E.J. Lin, D.C. Herman, A.M. Questell, S.E. Martello, R.J. Hedges, A.J. Walker, M. Rafat, Irradiated mammary spheroids elucidate Mechanisms of macrophage-mediated breast cancer recurrence, Cell. Mol. Bioeng. (2023). 10.1007/s12195-023-00775-x.

[8] J. Lou, D.J. Mooney, Chemical strategies to engineer hydrogels for cell culture, Nat. Rev. Chem. 6 (2022) 726–744. 10.1038/s41570-022-00420-7.

[9] H. Jin, Y. Yoo, Y. Kim, Y. Kim, J. Cho, Y.S. Lee, Radiation-induced lung fibrosis: Preclinical animal models and therapeutic strategies, Cancers (Basel). 12 (2020) 1–24. 10.3390/cancers12061561.

[10] M. Blatchley, J.S. Bader, A. Pandey, D. Pardoll, Tissue matrix arrays for high throughput screening and systems analysis of cell function, 12 (2016) 1197–1204. https://www.nature.com/articles/nmeth.3619.

[11] E. Henke, R. Nandigama, S. Ergün, Extracellular matrix in the tumor microenvironment and its impact on cancer therapy, Front. Mol. Biosci. 6 (2020) 1–24. 10.3389/fmolb.2019.00160.

[12] M.T. Wolf, K.A. Daly, E.P. Brennan-Pierce, S.A. Johnson, C.A. Carruthers, A. D’Amore, S.P. Nagarkar, S.S. Velankar, S.F. Badylak, A hydrogel derived from decellularized dermal extracellular matrix, Biomaterials. 33 (2012) 7028–7038. 10.1016/j.biomaterials.2012.06.051.

[13] L.E. Flynn, The use of decellularized adipose tissue to provide an inductive microenvironment for the adipogenic differentiation of human adipose-derived stem cells, Biomaterials. 31 (2010) 4715–4724. 10.1016/j.biomaterials.2010.02.046.

[14] M.T. Spang, K.L. Christman, Extracellular matrix hydrogel therapies: In vivo applications and development, Acta Biomater. 68 (2018) 1–14. 10.1016/j.actbio.2017.12.019.

[15] S.M. Alves, T. Zhu, A. Shostak, N.S. Rossen, M. Rafat, Studying normal tissue radiation effects using extracellular matrix hydrogels, J. Vis. Exp. (2019). 10.3791/59304.

[16] W. Ying, P.S. Cheruku, F.W. Bazer, S.H. Safe, B. Zhou, Investigation of macrophage polarization using bone marrow derived macrophages, J. Vis. Exp. (2013) 1–8. 10.3791/50323.

[17] J. Weischenfeldt, B. Porse, Bone marrow-derived macrophages (BMM): Isolation and applications, Cold Spring Harb. Protoc. 3 (2008) 1–7. 10.1101/pdb.prot5080.

[18] J. Schindelin, I. Arganda-Carreras, E. Frise, V. Kaynig, M. Longair, T. Pietzsch, S. Preibisch, C. Rueden, S. Saalfeld, B. Schmid, J.Y. Tinevez, D.J. White, V. Hartenstein, K. Eliceiri, P. Tomancak, A. Cardona, Fiji: An open-source platform for biological-image analysis, Nat. Methods. 9 (2012) 676–682. 10.1038/nmeth.2019.

[19] A. D’Amore, J.A. Stella, W.R. Wagner, M.S. Sacks, Characterization of the complete fiber network topology of planar fibrous tissues and scaffolds, Biomaterials. 31 (2010) 5345– 5354. 10.1016/j.biomaterials.2010.03.052.

[20] O. Chaudhuri, S.T. Koshy, C. Branco, J. Shin, C.S. Verbeke, K.H. Allison, D.J. Mooney, Extracellular matrix stiffness and composition jointly regulate the induction of malignant phenotypes in mammary epithelium, Nat. Mater. 13.10 (2014) 970–078. 10.1038/NMAT4009.

[21] M.J. Goldman, B. Craft, M. Hastie, K. Repecka, F. McDade, A. Kamath, A, Banerjee, Y. Luo, D. Rogers, A.N. Brooks, J. Zhu, D. Haussler, Visualizing and interpreting cancer genomics data via the Xena platform, Nat. Biotechnol. 38.6 (2020) 675–678. 10.1038/s41587-020-0546-8

[22] The Cancer Genome Atlas Network, Comprehensive molecular portraits of human breast tumours, Nature. 490.7418 (2012) 61–70. 10.1038/nature11412.

[23] J. Depciuch, E. Kaznowska, I. Zawlik, R. Wojnarowska, M. Cholewa, P. Heraud, J. Cebulski, Application of Raman spectroscopy and infrared spectroscopy in the identification of breast cancer, Appl. Spectrosc. 70 (2016) 251–263. 10.1177/0003702815620127.

[24] Z. Movasaghi, S. Rehman, I.U. Rehman, Raman spectroscopy of biological tissues, Appl. Spectrosc. Rev. 42 (2007) 493–541. 10.1080/05704920701551530.

[25] C. Gullekson, L. Lucas, K. Hewitt, L. Kreplak, Surface-Sensitive Raman Spectroscopy of Collagen I Fibrils Tip preparation, Biophysj. 100 (2011) 1837–1845. 10.1016/j.bpj.2011.02.026.

[26] H. Zhang, A.C. Silva, W. Zhang, H. Rutigliano, A. Zhou, Raman spectroscopy characterization extracellular vesicles from bovine placenta and peripheral blood mononuclear cells, PLoS One 15.7 (2020) e0235214. 10.1371/journal.pone.0235214.

[27] R. Ahmed, A.W.L. Law, T.W. Cheung, C. Lau, Raman spectroscopy of bone composition during healing of subcritical calvarial defects, Biomed. Opt. Express. 9 (2018) 1704. 10.1364/boe.9.001704.

[28] Y. Ling, H.F. Rios, E.R. Myers, Y. Lu, J.Q. Feng, A.L. Boskey, DMP1 depletion decreases bone mineralization in vivo: An FTIR imaging analysis, J. Bone Miner. Res. 20 (2005) 2169–2177. 10.1359/JBMR.050815.

[29] N. Kuhar, S. Sil, T. Verma, S. Umapathy, Challenges in application of Raman spectroscopy to biology and materials, RSC Adv. 8 (2018) 25888–25908. 10.1039/c8ra04491k.

[30] R.H.M. van der Meijden, D. Daviran, L. Rutten, X.F. Walboomers, E. Macías-Sánchez, N. Sommerdijk, A. Akiva, A 3D Cell-free Bone model shows collagen mineralization is driven and controlled by the Matrix, Adv. Funct. Mater. 33 (2023) 1–12. 10.1002/adfm.202212339.

[31] S.K. Paidi, A. Rizwan, C. Zheng, M. Cheng, K. Glunde, I. Barman, Label-free raman spectroscopy detects stromal adaptations in premetastatic lungs primed by breast cancer, Cancer Res. (2017). 10.1158/0008-5472.CAN-16-1862.

[32] J. Lever, M. Krzywinski, N. Altman, Points of Significance: Principal component analysis, Nat. Methods. 14 (2017) 641–642. 10.1038/nmeth.4346.

[33] X. Wen, Y.C. Ou, G. Bogatcheva, G. Thomas, A. Mahadevan-Jansen, B. Singh, E.C. Lin, R. Bardhan, Probing metabolic alterations in breast cancer in response to molecular inhibitors with Raman spectroscopy and validated with mass spectrometry, Chem. Sci. 11 (2020) 9863–9874. 10.1039/d0sc02221g.

[34] F.M. Lartey, M. Rafat, M. Negahdar, A. V. Malkovskiy, X. Dong, X. Sun, M. Li, T. Doyle, J. Rajadas, E.E. Graves, B.W. Loo, P.G. Maxim, Dynamic CT imaging of volumetric changes in pulmonary nodules correlates with physical measurements of stiffness, Radiother. Oncol. 122 (2017) 313–318. 10.1016/j.radonc.2016.11.019.

[35] F. Revert, F. Revert-Ros, R. Blasco, A. Artigot, E. López-Pascual, R. Gozalbo-Rovira, I. Ventura, E. Gutiérrez-Carbonell, N. Roda, D. Ruíz-Sanchis, J. Forteza, J. Alcácer, A. Pérez-Sastre, A. Díaz, E. Pérez-Payá, J.F. Sanz-Cervera, J. Saus, Selective targeting of collagen IV in the cancer cell microenvironment reduces tumor burden, Oncotarget. 9 (2018) 11020–11045. 10.18632/oncotarget.24280.

[36] H. JingSong, G. Hong, J. Yang, Z. Duo, F. Li, C. WeiCai, L. XueYing, M. YouSheng, O.Y. YiWen, P. Yue, C. Zou, siRNA-Mediated suppression of collagen type iv alpha 2 (COL4A2) mRNA inhibits triple-negative breast cancer cell proliferation and migration, Oncotarget. 8 (2017) 2585–2593. 10.18632/oncotarget.13716.

[37] S.P. Reese, N. Farhang, R. Poulson, G. Parkman, J.A. Weiss, Nanoscale imaging of collagen gels with focused ion beam milling and scanning electron microscopy, Biophys. J. 111 (2016) 1797–1804. 10.1016/j.bpj.2016.08.039.

[38] L. Yuan, B. Li, J. Yang, Y. Ni, Y. Teng, L. Guo, H. Fan, Y. Fan, X. Zhang, Effects of composition and mechanical property of injectable collagen I/II composite hydrogels on chondrocyte behaviors, Tissue Eng. - Part A. 22 (2016) 899–906. 10.1089/ten.tea.2015.0513.

[39] R. Zhao, A. Afthinos, T. Zhu, P. Mistriotis, Y. Li, S.A. Serra, Y. Zhang, C.L. Yankaskas, S. He, M.A. Valverde, S.X. Sun, K. Konstantopoulos, Cell sensing and decision-making in confinement: The role of TRPM7 in a tug of war between hydraulic pressure and cross-sectional area, Sci. Adv. 5 (2019). 10.1126/sciadv.aaw7243.

[40] J. Stricker, T. Falzone, M.L. Gardel, Mechanics of the F-actin cytoskeleton, J. Biomech. (2010). 10.1016/j.jbiomech.2009.09.003.

[41] P.A. Janmey, C.A. McCulloch, Cell mechanics: Integrating cell responses to mechanical stimuli, Annu. Rev. Biomed. Eng. 9 (2007) 1–34. 10.1146/annurev.bioeng.9.060906.151927.

[42] B. Emon, J. Bauer, Y. Jain, B. Jung, T. Saif, Biophysics of tumor microenvironment and cancer metastasis - A mini review, Comput. Struct. Biotechnol. J. 16 (2018) 279–287. 10.1016/j.csbj.2018.07.003.

[43] M.P. Iwanicki, T. Vomastek, R.W. Tilghman, K.H. Martin, J. Banerjee, P.B. Wedegaertner, J.T. Parsons, FAK, PDZ-RhoGEF and ROCKII cooperate to regulate adhesion movement and trailing-edge retraction in fibroblasts, J. Cell Sci. 121 (2008) 895–905. 10.1242/jcs.020941.

[44] H.S. Carr, Y. Zuo, W. Oh, J.A. Frost, Regulation of focal adhesion kinase activation, breast cancer cell motility, and amoeboid invasion by the RhoA guanine nucleotide exchange factor net1, Mol. Cell. Biol. 33 (2013) 2773–2786. 10.1128/mcb.00175-13.

[45] H. Paz, N. Pathak, J. Yang, Invading one step at a time: The role of invadopodia in tumor metastasis, Oncogene. 33 (2014) 4193–4202. 10.1038/onc.2013.393.

[46] G. Bati, D. Pesen Okvur, Invadopodia: Proteolytic feet of cancer cells, Turkish J. Biol. 38 (2014) 740–747. 10.3906/biy-1404-110.

[47] M.J. Breiding, Matrix rigidity differentially regulates invadopodia activity through ROCK1 and ROCK2, Physiol. Behav. 63 (2014) 1–18. 10.1016/j.biomaterials.2016.01.028.

[48] M. Noi, K.I. Mukaisho, S. Yoshida, S. Murakami, S. Koshinuma, T. Adachi, Y. Machida, M. Yamori, T. Nakayama, G. Yamamoto, H. Sugihara, ERK phosphorylation functions in invadopodia formation in tongue cancer cells in a novel silicate fibre-based 3D cell culture system, Int. J. Oral Sci. 10 (2018) 1–10. 10.1038/s41368-018-0033-y.

[49] H. Yamaguchi, M. Lorenz, S. Kempiak, C. Sarmiento, S. Coniglio, M. Symons, J. Segall, R. Eddy, H. Miki, T. Takenawa, J. Condeelis, Molecular mechanisms of invadopodium formation: The role of the N-WASP-Arp2/3 complex pathway and cofilin, J. Cell Biol. 168 (2005) 441–452. 10.1083/jcb.200407076.

[50] J. Nam, M. Kang, A.M. Suchar, T. Shimamura, E.A. Kohn, A.M. Michalowska, V.C. Jordan, S. Hirohashi, M. Lalage, NIH Public Access, 66 (2007) 7176–7184.

[51] M. Raškov, A. Venhauerov, M. Jakubek, D. Rosel, K. Smetana, The Role of IL-6 in cancer cell invasiveness and metastasis — overview and therapeutic opportunities, Cells. 11.22 (2022) 3698. 10.3390/cells11223698

[52] X. Sun, D.J. Glynn, L.J. Hodson, C. Huo, K. Britt, E.W. Thompson, L. Woolford, A. Evdokiou, J.W. Pollard, S.A. Robertson, W. V. Ingman, CCL2-driven inflammation increases mammary gland stromal density and cancer susceptibility in a transgenic mouse model, Breast Cancer Res. 19 (2017) 1–15. 10.1186/s13058-016-0796-z.

[53] J. Wang, D. Li, H. Cang, B. Guo, Crosstalk between cancer and immune cells: Role of tumor-associated macrophages in the tumor microenvironment, Cancer Med. 8 (2019) 4709–4721. 10.1002/cam4.2327.

[54] T. Ryszer, Understanding the mysterious M2 macrophage through activation markers and effector mechanisms, Mediators Inflamm. (2015) 16–18. 10.1155/2015/816460.

[55] S. Eshghjoo, D.M. Kim, A. Jayaraman, Y. Sun, R.C. Alaniz, Macrophage polarization in atherosclerosis, Genes (Basel). 13 (2022) 1–10. 10.3390/genes13050756.

[56] Y. Yao, X.H. Xu, L. Jin, Macrophage polarization in physiological and pathological pregnancy, Front. Immunol. 10 (2019) 1–13. 10.3389/fimmu.2019.00792.

[57] I. Tvaroška, S. Kozmon, J. Kóňa, Molecular modeling insights into the structure and behavior of integrins: A review, Cells. 12 (2023) 1–44. 10.3390/cells12020324.

[58] J. Cooper, F.G. Giancotti, Integrin signaling in cancer: mechanotransduction, stemness, epithelial plasticity, and therapeutic resistance, Cancer Cell. 35 (2019) 347–367. 10.1016/j.ccell.2019.01.007.

[59] Y. Shu, M. Qin, Y. Song, Q. Tang, Y. Huang, P. Shen, Y. Lu, M2 polarization of tumor-associated macrophages is dependent on integrin β3 via peroxisome proliferator-activated receptor-γ up-regulation in breast cancer, Immunology. 160 (2020) 345–356. 10.1111/imm.13196.

[60] L. Zhang, Y. Dong, Y. Dong, J. Cheng, J. Du, Role of integrin-β3 protein in macrophage polarization and regeneration of injured muscle, J. Biol. Chem. 287 (2012) 6177–6186. 10.1074/jbc.M111.292649.

[61] M.R. A.D. Sherry, R. Eyben, N.B. Newman, P. Gutkin, I. Mayer, K. Horst, A. B. Chakravarthy, Systemic inflammation after radiation predicts locoregional recurrence, progression, and mortality in stage II-III Triple-Negative Breast Cancer, Int J Radiat Oncol Biol Phys. 108 (2020) 268–276. 10.1016/j.ijrobp.2019.11.398.

[62] K.M. Arnold, N.J. Flynn, A. Raben, L. Romak, Y. Yu, A.P. Dicker, F. Mourtada, J. Sims-Mourtada, The impact of radiation on the tumor microenvironment: effect of dose and fractionation schedules, Cancer Growth Metastasis. 11 (2018) 117906441876163. 10.1177/1179064418761639.

[63] J.M. Straub, J. New, C.D. Hamilton, C. Lominska, S.M. Thomas, Radiation-induced fibrosis: mechanisms and implications for therapy, Radiation-Induced Fibros. J. Cancer. Res. Clin. Oncol.141 (2016) 1–16. 10.1007/s00432-015-1974-6.Radiation-induced.

[64] T.J. Bledsoe, S.K. Nath, R.H. Decker, Radiation pneumonitis, Clin. Chest Med. 38.2 (2017) 201–208. http://doi/org/10.1016/j.ccm.2016.12.004.

[65] M.J. Breiding, Tumor-associated fibrosis as a regulator of tumor immunity and response to immunotherapy, Physiol. Behav. 63 (2014) 1–18. 10.1007/s00262-017-2003-1.Tumor-associated.

[66] V. Poltavets, M. Kochetkova, S.M. Pitson, M.S. Samuel, The role of the extracellular matrix and its molecular and cellular regulators in cancer cell plasticity, Front. Oncol. 8 (2018) 1–19. 10.3389/fonc.2018.00431.

[67] P. Lu, V.M. Weaver, Z. Werb, The extracellular matrix: A dynamic niche in cancer progression, J. Cell Biol. 196 (2012) 395–406. 10.1083/jcb.201102147.

[68] L.A. Northcutt1, A. Suarez-Arnedo, M. Rafat, Emerging biomimetic materials for studying tumor and immune cell behavior, Ann Biomed Eng. 48 (2020) 2064–2077. 10.1007/s10439-019-02384-0.

[69] C. Herskind, C. Sticht, A. Sami, F.A. Giordano, F. Wenz, Gene expression profiles reveal extracellular matrix and inflammatory signaling in radiation-induced premature differentiation of human fibroblast in vitro, Front. Cell Dev. Biol. 9 (2021) 1–17. 10.3389/fcell.2021.539893.

[70] L. Shukla, R. Luwor, M.E. Ritchie, S. Akabarzadeh, H.J. Zhu, W. Morrison,T. Karnezis, R. Shayan, Therapeutic reversal of radiotherapy injury to pro-fibrotic dysfunctional fibroblasts in vitro using adipose-derived stem cells, PRS Global Open 8.3 (2020). 10.1097/GOX.0000000000002706.

[71] Z. Wang, Y. Tang, Y. Tan, Q. Wei, W. Yu, Cancer-associated fibroblasts in radiotherapy: Challenges and new opportunities, Cell Commun. Signal. 17 (2019) 1–12. 10.1186/s12964-019-0362-2.

[72] G. Bahcecioglu, X. Yue, E. Howe, I. Guldner, M.S. Stack, H. Nakshatri, S. Zhang, P. Zorlutuna, Aged breast extracellular matrix Drives mammary epithelial cells to an invasive and cancer-like phenotype, Adv. Sci. 8 (2021) 1–15. 10.1002/advs.202100128.

[73] K. Song, Z. Yu, X. Zu, G. Li, Z. Hu, Y. Xue, Collagen remodeling along cancer progression providing a novel opportunity for cancer diagnosis and treatment, Int. J. Mol. Sci. 23 (2022). 10.3390/ijms231810509.

[74] B. Wang, J. Wei, L. Meng, H. Wang, C. Qu, X. Chen, Y. Xin, X. Jiang, Advances in pathogenic mechanisms and management of radiation-induced fibrosis, Biomed. Pharmacother. 121 (2020) 109560. 10.1016/j.biopha.2019.109560.

[75] Y. Deng, P. Chakraborty, M.K. Jolly, H. Levine, A theoretical approach to coupling the epithelial-mesenchymal transition (EMT) to extracellular matrix (ECM) stiffness via loxl2, Cancers (Basel). 13 (2021) 1–14. 10.3390/cancers13071609.

[76] O. Maller, A.P. Drain, A.S. Barrett, S. Borgquist, B. Ruffell, I. Zakharevich, T.T. Pham, T. Gruosso, H. Kuasne, J.N. Lakins, I. Acerbi, J.M. Barnes, T. Nemkov, A. Chauhan, J. Gruenberg, A. Nasir, O. Bjarnadottir, Z. Werb, P. Kabos, Y.Y. Chen, E.S. Hwang, M. Park, L.M. Coussens, A.C. Nelson, K.C. Hansen, V.M. Weaver, Tumour-associated macrophages drive stromal cell-dependent collagen crosslinking and stiffening to promote breast cancer aggression, Nat. Mater. 20 (2021) 548–559. 10.1038/s41563-020-00849-5.

[77] L.D. Muiznieks, F.W. Keeley, Molecular assembly and mechanical properties of the extracellular matrix: A fibrous protein perspective, Biochim. Biophys. Acta - Mol. Basis Dis. 1832 (2013) 866–875. 10.1016/j.bbadis.2012.11.022.

[78] B. Yue, Biology of the extracellular matrix: an overview, J. Glaucoma. (2014) S20–S23. 10.1097/IJG.0000000000000108.

[79] J.P. Fatherree, J.R. Guarin, R.A. McGinn, S.P. Naber, M.J. Oudin, Chemotherapy - induced collagen IV drives cancer cell motility through activation of Src and focal adhesion kinase, Cancer. Res. 82.10 (2022) 2031–2044. 10.1158/0008-5472.CAN-21-1823

[80] P. Iyengar, V. Espina, T.W. Williams, Y. Lin, D. Berry, L.A. Jelicks, H. Lee, K. Temple, R. Graves, J. Pollard, N. Chopra, R.G. Russell, R. Sasisekharan, B.J. Trock, M. Lippman, V.S. Calvert, E.F. Petricoin, L. Liotta, E. Dadachova, R.G. Pestell, M.P. Lisanti, P. Bonaldo, P.E. Scherer, Adipocyte-derived collagen VI affects early mammary tumor progression in vivo, demonstrating a critical interaction in the tumor/stroma microenvironment, J. Clin. Invest. 115 (2005) 1163–1176. 10.1172/JCI23424.

[81] A.L. Wishart, S.J. Conner, J.R. Guarin, J.P. Fatherree, Y. Peng, R.A. McGinn, R. Crews, S.P. Naber, M. Hunter, A.S. Greenberg, M.J. Oudin, Decellularized extracellular matrix scaffolds identify full-length collagen VI as a driver of breast cancer cell invasion in obesity and metastasis, Sci. Adv. 6 (2020). 10.1126/sciadv.abc3175.

[82] D.M. Gilkes, G.L. Semenza, D. Wirtz, Hypoxia and the extracellular matrix: Drivers of tumour metastasis, Nat. Rev. Cancer. 14 (2014) 430–439. 10.1038/nrc3726.

[83] M. Ansems, P.N. Span, The tumor microenvironment and radiotherapy response; a central role for cancer-associated fibroblasts, Clin. Transl. Radiat. Oncol. 22 (2020) 90–97. 10.1016/j.ctro.2020.04.001.

[84] W.H. Lee, J.P. Warrington, W.E. Sonntag, Irradiation alters MMP-2/TIMP-2 system and collagen type IV degradation in brain, IJROBP, 82.5 (2012) 1559–1566. 10.1016/j.ijrobp.2010.12.032

[85] R.N. Andrews, D.L. Caudell, L.J. Metheny-Barlow, A.M. Peiffer, J.A. Tooze, J.D. Bourland, R.E. Hampson, S.A. Deadwyler, J.M. Cline, Fibronectin produced by cerebral endothelial and vascular smooth muscle cells contributes to perivascular extracellular matrix in late-delayed radiation-induced brain injury, Radiat. Res. 190 (2018) 361–373. 10.1667/RR14961.1.

[86] F. Mohamed, D.A. Bradley, C.P. Winlove, Effects of ionizing radiation on extracellular matrix, Nucl. Instruments Methods Phys. Res. Sect. A Accel. Spectrometers, Detect. Assoc. Equip. 580 (2007) 566–569. 10.1016/J.NIMA.2007.05.236.

[87] R.A. Stile, K.E. Healy, Thermo-responsive peptide-modified hydrogels for tissue regeneration, Biomacromolecules. 2 (2001) 185–194. 10.1021/bm0000945.

[88] M.W. Tibbitt, K.S. Anseth, Hydrogels as extracellular matrix mimics for 3D cell culture, Biotechnol. Bioeng. 103 (2009) 655–663. 10.1002/bit.22361.

[89] C. Matthew T. Wolfa, b, Kerry A. Dalyb, c, Janet E. Reingb, and Stephen F. Badylaka, b, Biologic scaffold composed of skeletal muscle extracellular matrix, Physiol. Behav. 63 (2014) 1–18. 10.1016/j.biomaterials.2011.12.055.Biologic.

[90] D.A. Young, D.O. Ibrahim, D. Hu, K.L. Christman, Injectable hydrogel scaffold from decellularized human lipoaspirate, Acta Biomater. 7 (2011) 1040–1049. 10.1016/j.actbio.2010.09.035.

[91] T.L. Sellaro, D. Ph, A. Ranade, D. Ph, D.M. Faulk, G.P. Mccabe, D. Ph, K. Dorko, S.F. Badylak, D. Ph, S.C. Strom, D. Ph, Maintenance of Human Hepatocyte Function In Vitro by Liver-Derived Extracellular Matrix Gels, Tissue Eng. Part A. 16 (2010).

[92] J. Zhou, P. Wu, H. Sun, H. Zhou, Y. Zhang, Z. Xiao, Lung tissue extracellular matrix-derived hydrogels protect against radiation-induced lung injury by suppressing epithelial– mesenchymal transition, J. Cell. Physiol. 235 (2020) 2377–2388. 10.1002/jcp.29143.

[93] R.H.J. De Hilster, P.K. Sharma, M.R. Jonker, E.S. White, E.A. Gercama, M. Roobeek, W. Timens, M.C. Harmsen, M.N. Hylkema, J.K. Burgess, Human lung extracellular matrix hydrogels resemble the stiffness and viscoelasticity of native lung tissue, Am. J. Physiol. - Lung Cell. Mol. Physiol. 318 (2020) L698–L704. 10.1152/AJPLUNG.00451.2019.

[94] K.F. Ruud, W.C. Hiscox, I. Yu, R.K. Chen, W. Li, Distinct phenotypes of cancer cells on tissue matrix gel, Breast Cancer Res. 22 (2020) 1–22. 10.1186/s13058-020-01321-7.

[95] L. Cassereau, Y.A. Miroshnikova, G. Ou, J. Lakins, V.M. Weaver, A 3D tension bioreactor platform to study the interplay between ECM stiffness and tumor phenotype, J. Biotechnol. 193 (2015) 66–69. 10.1016/j.jbiotec.2014.11.008.

[96] P.L. Chandran, V.H. Barocas, Deterministic material-based averaging theory model of collagen gel micromechanics, J. Biomech. Eng. 129 (2007) 137–147. 10.1115/1.2472369.

[97] J. Zonderland, L. Moroni, Steering cell behavior through mechanobiology in 3D: A regenerative medicine perspective, Biomaterials. 268 (2021) 120572. 10.1016/j.biomaterials.2020.120572.

[98] V.Z. Beachley, M.T. Wolf, K. Sadtler, S.S. Manda, H. Jacobs, M.R. Blatchley, J.S. Bader, A. Pandey, D. Pardoll, J.H. Elisseeff, Tissue matrix arrays for high-throughput screening and systems analysis of cell function, Nat. Methods. (2015). 10.1038/nmeth.3619.

[99] N. Cordes, Integrin-mediated cell-matrix interactions for prosurvivaland antiapoptotic signaling after genotoxic injury, Cancer Lett. 242 (2006) 11–19. 10.1016/j.canlet.2005.12.004.

[100] J.W. Denham, M. Hauer-Jensen, The radiotherapeutic injury - A complex “wound,” Radiother. Oncol. 63 (2002) 129–145. 10.1016/S0167-8140(02)00060-9.

[101] C. Medrek, F. Pontén, K. Jirström, K. Leandersson, The presence of tumor associated macrophages in tumor stroma as a prognostic marker for breast cancer patients, BMC Cancer. 12 (2012) 1. 10.1186/1471-2407-12-306.

[102] S.K. Biswas, A. Mantovani, Macrophage plasticity and interaction with lymphocyte subsets: Cancer as a paradigm, Nat. Immunol. 11 (2010) 889–896. 10.1038/ni.1937.

[103] A. Yuan, Y.J. Hsiao, H.Y. Chen, H.W. Chen, C.C. Ho, Y.Y. Chen, Y.C. Liu, T.H. Hong, S.L. Yu, J.J.W. Chen, P.C. Yang, Opposite effects of M1 and M2 macrophage subtypes on lung cancer progression, Sci. Rep. 5 (2015) 1–12. 10.1038/srep14273.

[104] E.S. Stovgaard, D. Nielsen, E. Hogdall, E. Balslev, Triple negative breast cancer– prognostic role of immune-related factors: a systematic review, Acta Oncol. (Madr). 57 (2018) 74–82. 10.1080/0284186X.2017.1400180.

[105] I. Shabo, O. Stål, H. Olsson, S. Doré, J. Svanvik, Breast cancer expression of CD163, a macrophage scavenger receptor, is related to early distant recurrence and reduced patient survival, Int. J. Cancer. 123 (2008) 780–786. 10.1002/ijc.23527.

[106] Y. Xue, L. Tong, F. Liu, A. Liu, S. Zeng, Q. Xiong, Z. Yang, X. He, Y. Sun, C. Xu, Tumor-infiltrating M2 macrophages driven by specific genomic alterations are associated with prognosis in bladder cancer, Oncol. Rep. 42 (2019) 581–594. 10.3892/or.2019.7196.

[107] D.G. DeNardo, D.J. Brennan, E. Rexhepaj, B. Ruffell, S.L. Shiao, S.F. Madden, W.M. Gallagher, N. Wadhwani, S.D. Keil, S.A. Junaid, H.S. Rugo, E. Shelley Hwang, K. Jirström, B.L. West, L.M. Coussens, Leukocyte complexity predicts breast cancer survival and functionally regulates response to chemotherapy, Cancer Discov. 1 (2011) 54–67. 10.1158/2159-8274.CD-10-0028.

[108] C. Margadant, H.N. Monsuur, J.C. Norman, A. Sonnenberg, Mechanisms of integrin activation and trafficking, Curr. Opin. Cell Biol. 23 (2011) 607–614. 10.1016/J.CEB.2011.08.005.

[109] R.O. Hynes, Integrins: bidirectional, allosteric signaling machines, Cell. 110.6 (2002) 673–687.10.1016/S0092-8674(02)00971-6

[110] T. Bui, J. Rennhack, S. Mok, C. Ling, M. Perez, J. Roccamo, E.R. Andrechek, C. Moraes, W.J. Muller, Functional Redundancy between β1 and β3 Integrin in activating the IR/Akt/mTORC1 Signaling Axis to Promote ErbB2-Driven Breast Cancer, Cell Rep. 29 (2019) 589–602.e6. 10.1016/j.celrep.2019.09.004.

