## Supplementary Figures for "Mammary Tissue-Derived Extracellular Matrix Hydrogels Reveal the Role of Irradiation in Driving a Pro-Tumor and Immunosuppressive Microenvironment"

**Supplemental Figures**

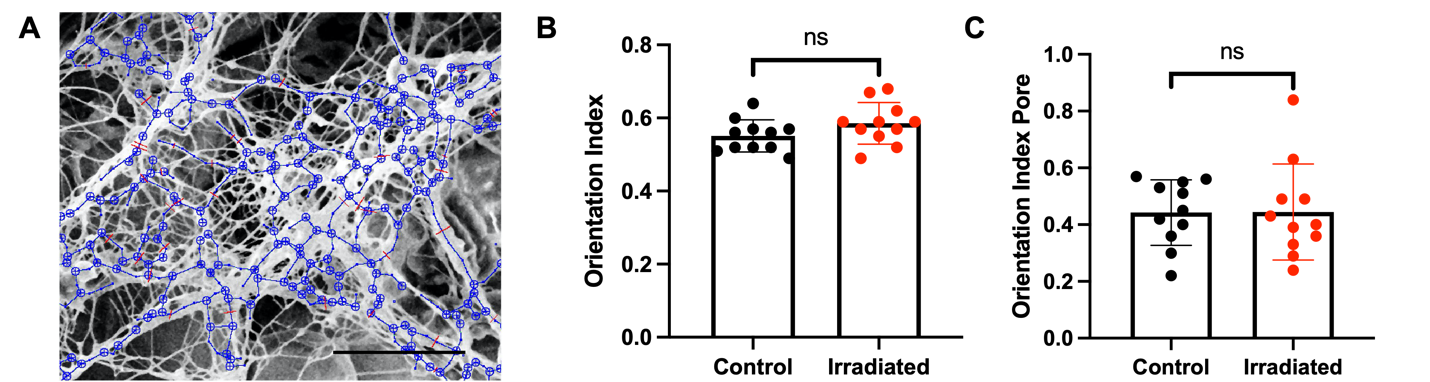

**Figure S1. Automated fiber tracking algorithm.** (**A**) Representative fiber network analysis on murine MFPs. Scale bar is 10 μm. SEM images were analyzed using an automated fiber tracking algorithm to determine the orientation index (**B**) and orientation index pore (**C**) of control and irradiated MFPs (n = 10). Error bars show standard deviation.

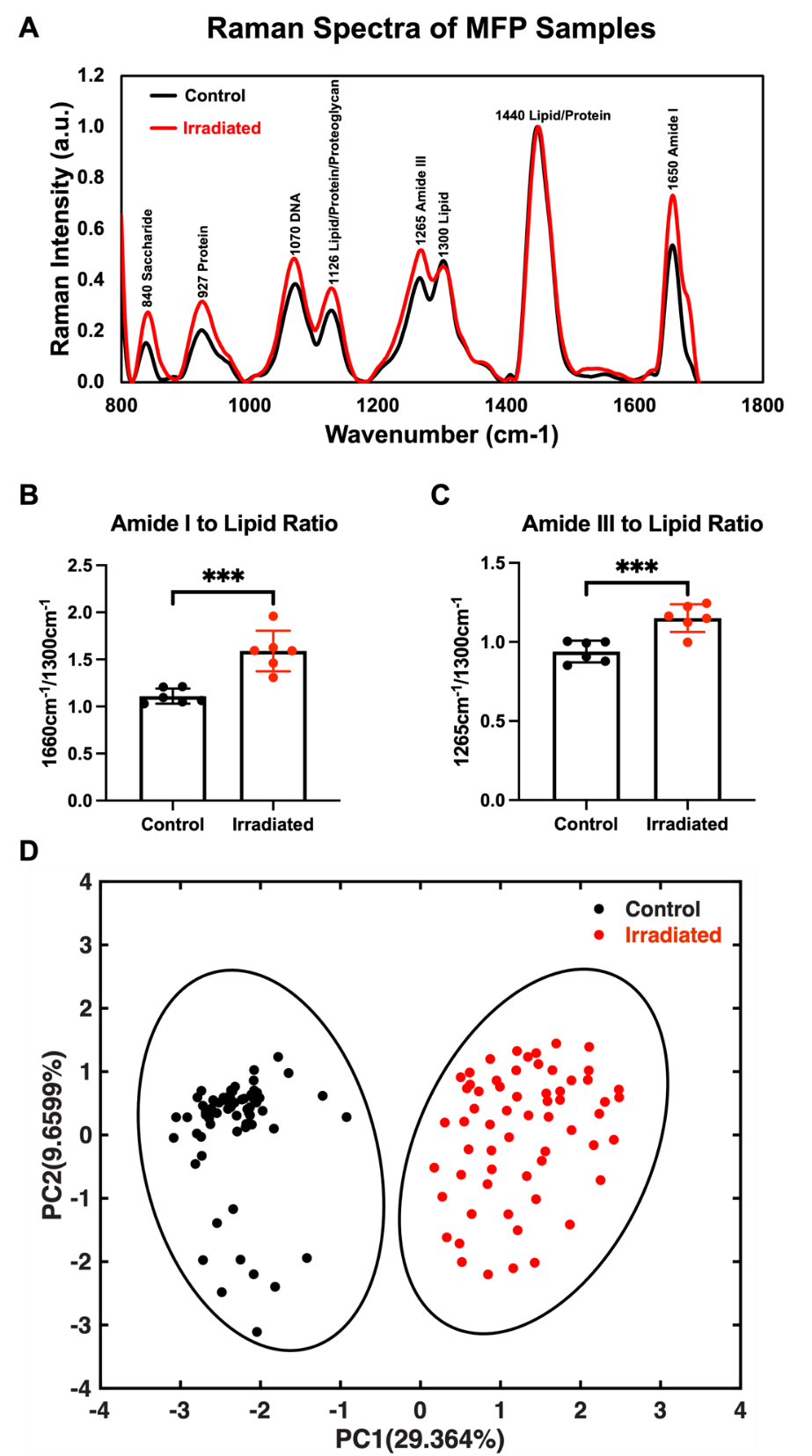

**Figure S2. Molecular composition of control and irradiated MFPs from Nu/Nu mice.** (**A**) Raman spectra of control and *ex vivo* irradiated (20 Gy) MFPs from Nu/Nu mice (n = 6). Spectra were normalized to the 1440 cm^-1^ biological peak. The peak intensity ratios of amide I to lipid (1660 cm^-1^/1300 cm^-1^) (**B**) and amide III to lipid (1265 cm^-1^/1300 cm^-1^) (**C**) are shown for control and irradiated samples. (**D**) PCA analysis of control and irradiated MFPs in 2D. Statistical significance was determined by Student’s t-test with ***p<0.001. Error bars show standard deviation.

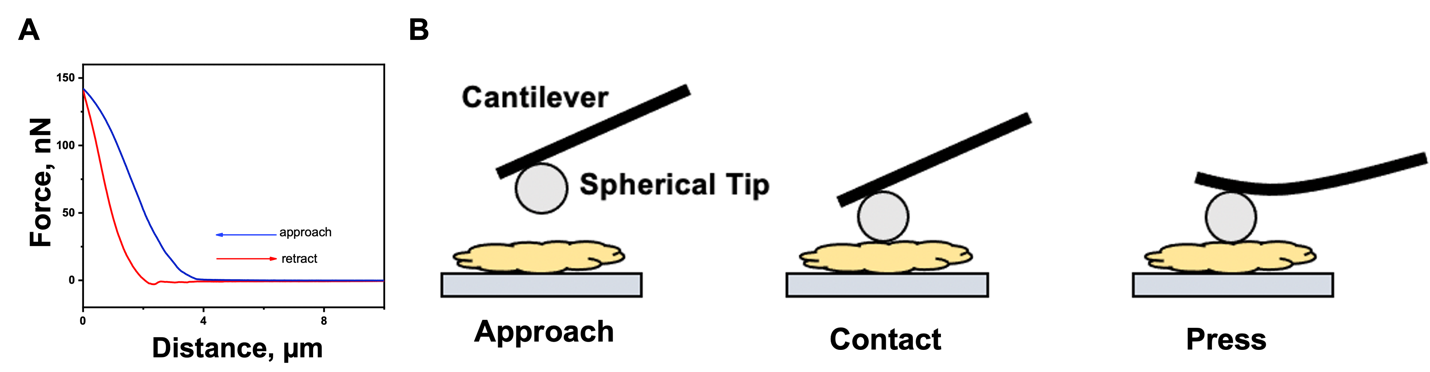

**Figure S3. Measuring the stiffness of MFPs using** [**atomic force microscopy**](https://www.sciencedirect.com/topics/medicine-and-dentistry/atomic-force-microscopy) **(AFM).** (**A**) Representative force-distance (FD) curve using the Hertz model to determine Young’s Modulus of MFPs. (**B**) Schematic of AFM using a spherical tip for FD measurements.

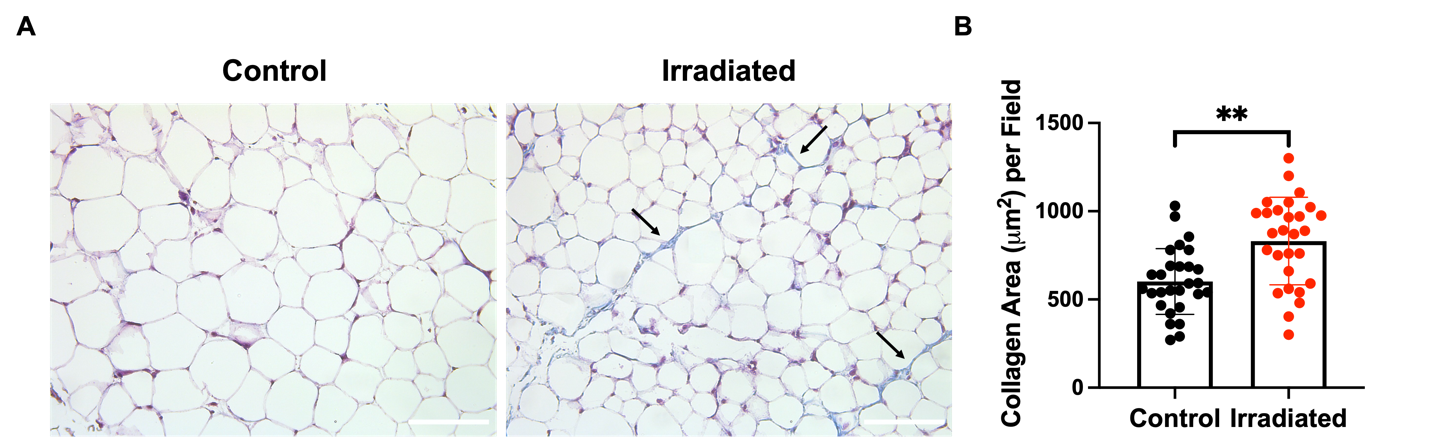

**Figure S4. Collagen content in control and irradiated MFPs.** Collagen in control and *ex vivo* irradiated MFPs was evaluated (**A**) and quantified (**B**) using Masson’s trichrome staining (n = 5). Arrows indicate positive staining. Scale bars are 100 μm. Statistical significance was determined by Student’s t-test with **p<0.01. Error bars show standard deviation.

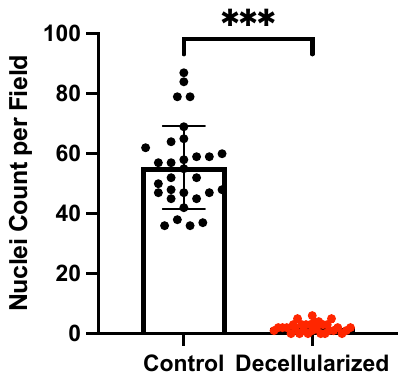

**Figure S5. Confirmation of nuclei removal in decellularized MFPs.** The number of nuclei per field in control and decellularized MFPs was quantified using H&E staining. Statistical significance was determined by Student’s t-test with **p<0.01 and ***p<0.001. Error bars show standard deviation.

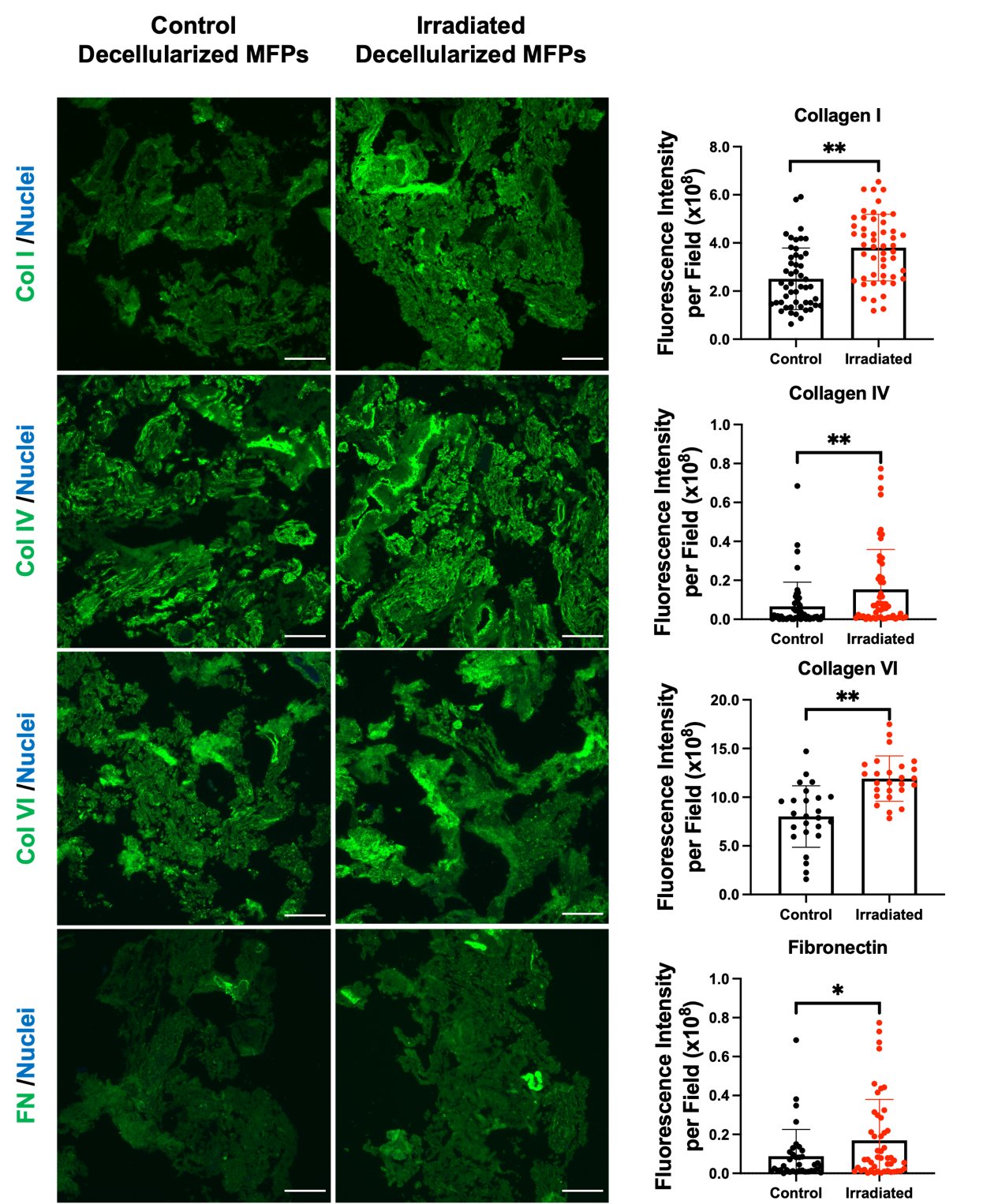

**Figure S6. Decellularized MFPs retain tissue ECM changes after radiation.** Immunofluorescence staining and quantification of collagen I (Col I), collagen IV (Col IV), collagen VI (Col VI), and fibronectin (FN) in decellularized control and irradiated MFPs per field (n = 4 mice). Scale bars are 100 μm. Statistical significance was determined by ANOVA analysis with *p<0.05 and **p<0.01. Error bars show standard deviation.

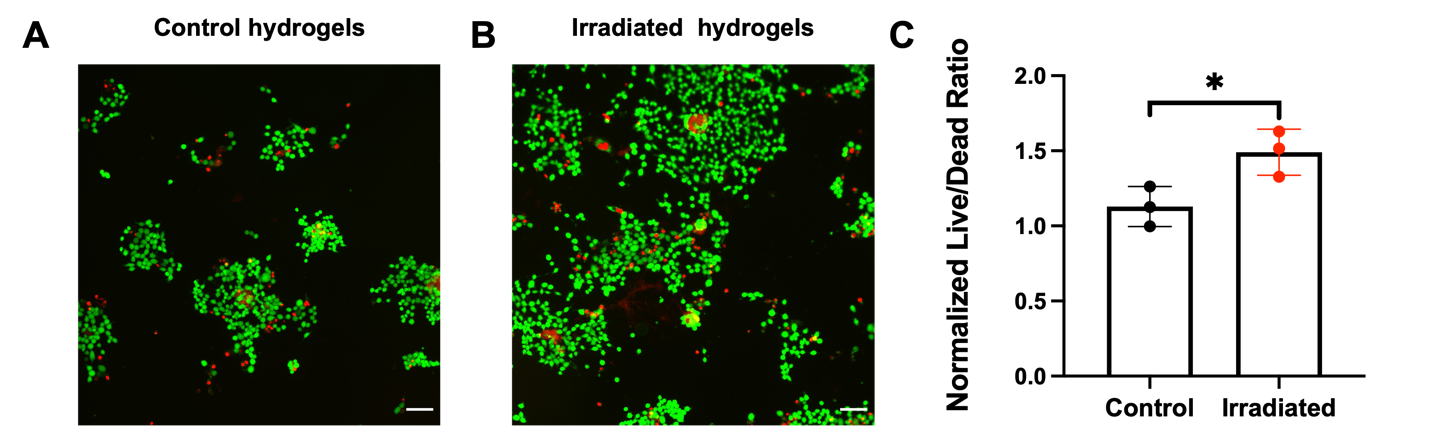

**Figure S7. Irradiated dECM hydrogels are not cytotoxic.** Fluorescence images of live (green) and dead (red) 4T1 cells in control (**A**) and *ex vivo* irradiated (**B**) dECM hydrogels from Nu/Nu mice seeded at 5x10^5^ cells/mL after 48 hr. (**C**) The fluorescence intensity ratio of live to dead cells encapsulated in ECM hydrogels (n = 3 biological replicates) was quantified and normalized to values from unirradiated controls. Scale bars are 100 μm. Statistical significance was determined by Student’s t-test with *p<0.05. Error bars show standard deviation.

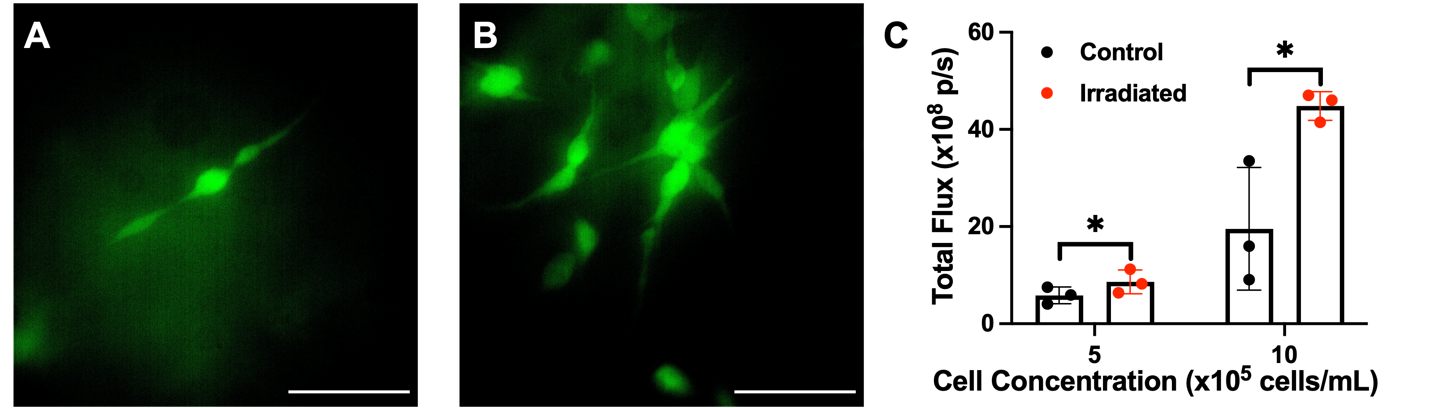

**Figure S8. MDA-MB-231 cell proliferation is enhanced in irradiated dECM hydrogels.** Fluorescence images of GFP- and luciferase-labeled human MDA-MB-231 TNBC cells in control (**A**) and *ex vivo* irradiated (**B**) dECM hydrogels from Nu/Nu mice (n = 3 per condition). Scale bars are 100 μm. (**C**) Bioluminescence quantification of MDA-MB-231 cells encapsulated in dECM hydrogels. Statistical significance was determined by Student’s t-test with *p<0.05. Error bars show standard deviation.

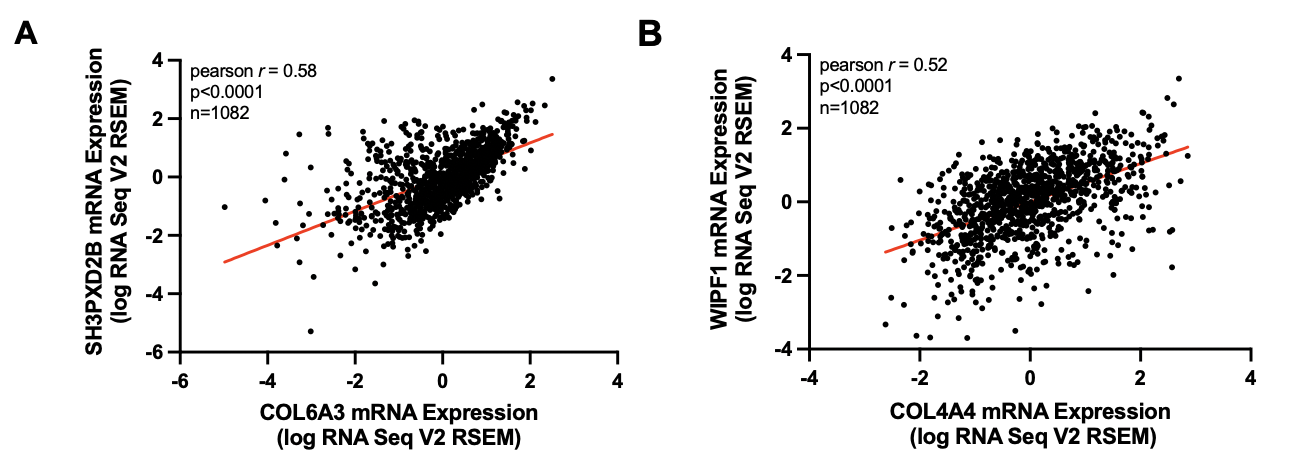

**Figure S9. Expression of collagen subtypes and invadopodia markers are correlated in human breast cancer.** Correlation of (**A**) *Col6A3* and *SH3PXD2B* mRNA expression and (**B**) *Col4A4* and *WIPF1* mRNA expression from TCGA analysis (PanCancer dataset) in 1,082 patients with breast tumors. Pearson correlation is shown.

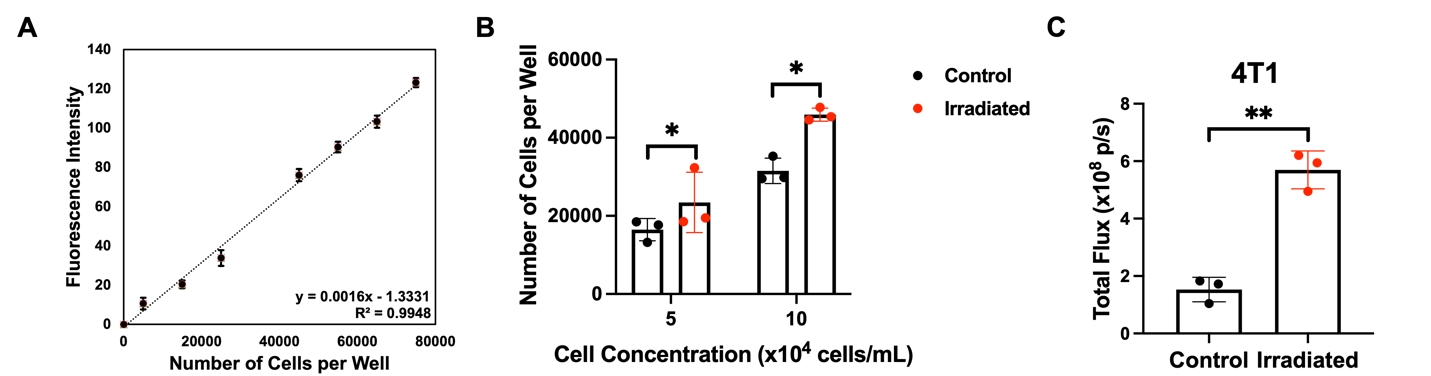

**Figure S10. Confirmation of BMDM and 4T1 proliferation in irradiated ECM microenvironments.** (**A**) Standard curve showing that alamarBlue signal is linear up to 80,000 BMDMs after a 4 hr incubation. (**B**) Quantified BMDM cell number in control and irradiated dECM hydrogels using the alamarBlue assay (n = 3 replicates). (**C**) Bioluminescence quantification of GFP- and luciferase-labeled 4T1 cells co-cultured with BMDMs (n = 3 replicates). Statistical significance was determined by Student’s t-test with *p<0.05. Error bars show the standard deviation.

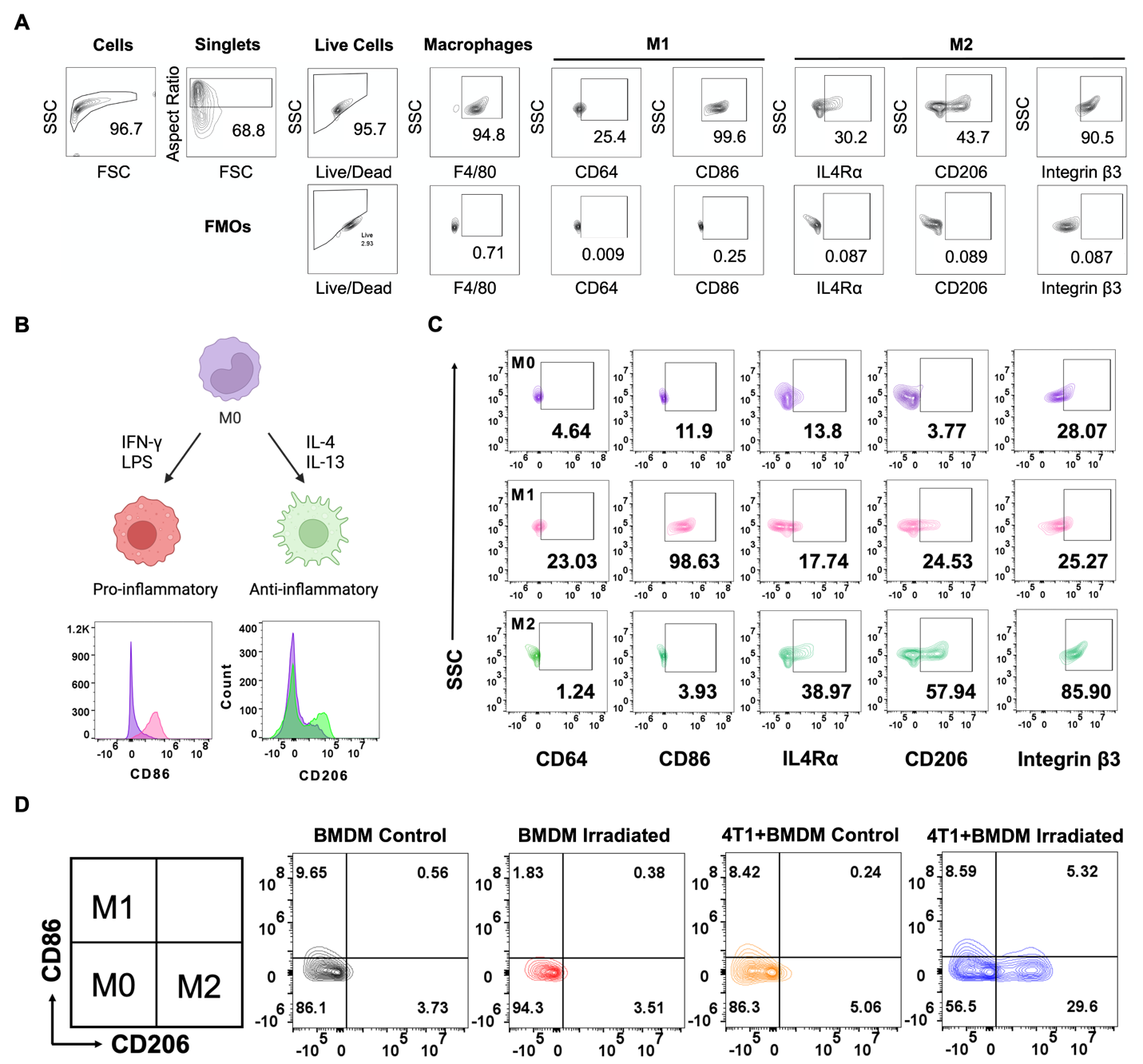

**Figure S11. Validation of macrophage polarization.** (**A**) Full gating strategy for macrophage markers with full minus one (FMO) controls shown in the bottom panels. (**B**) Scheme showing how M0 BMDMs were polarized to an M1 (interferon-gamma (IFN-γ), 100 µg/mL; lipopolysaccharide (LPS), 100 µg/mL for 24 hr) or M2 (interleukin-4 (IL-4), 100 µg/mL; IL-13, 100 µg/mL for 48 hr) phenotype with flow cytometry confirmation. (**C**) Flow cytometry gating of M0, M1, and M2 macrophages. (**D**) Expression of CD86 and CD206 in BMDMs isolated from individual-control, individual-irradiated, coculture-control, or coculture-irradiated dECM hydrogels. The percentage of positive cells per population of interest is reported.

**Tables**

**Table S1. List of cytokines evaluated using a Luminex 31-plex immunoassay.**

| **Fold Change > 1.5** | | |
| --- | --- | --- |
| **Name** | | **Abbreviation** |
| Granulocyte-macrophage colony-stimulating factor | | GM-CSF |
| Monocyte chemoattractant protein-1/  C-C Motif chemokine ligand 2 | | MCP-1/  CCL2 |
| Interleukin 6 | | IL-6 |
| Lipopolysaccharide-induced CXC chemokine/  Chemokine (C-X-C Motif) ligand 5 | | LIX/  CXCL5 |
| Leukemia inhibitory factor | | LIF |
| Monokine induced by gamma interferon/  Chemokine (C-X-C Motif) ligand 9 | | MIG/  CXCL9 |
| Interferon γ-induced protein 10 kDa/  Chemokine (C-X-C motif) ligand 10 | | IP-10/  CXCL10 |
| Tumor necrosis factor alpha | | TNFα |
| Interleukin 1 alpha | | IL-1α |
| **Fold Change < 1.5** | | |
| **Name** | **Abbreviation** | |
| Eotaxin | Eotaxin | |
| Granulocyte colony-stimulating factor | G-CSF | |
| Interferon gamma | IFNγ | |
| Interleukin 1 beta | IL-1β | |
| Interleukin 2 | IL-2 | |
| Interleukin 3 | IL-3 | |
| interleukin 4 | IL-4 | |
| Interleukin 5 | IL-5 | |
| Interleukin 7 | IL-7 | |
| Interleukin 9 | IL-9 | |
| Interleukin 10 | IL-10 | |
| Interleukin 12p40 | IL-12p40 | |
| Interleukin 12p70 | IL-12p70 | |
| Interleukin 13 | IL-13 | |
| Interleukin 15 | IL-15 | |
| Interleukin 17 | IL-17 | |
| Keratinocytes-derived chemokine | KC | |
| Macrophage colony-stimulating factor | M-CSF | |
| Macrophage inflammatory protein-1 alpha | MIP-1α | |
| Macrophage inflammatory protein-1 beta | MIP-1β | |
| Macrophage inflammatory protein-2 | MIP-2 | |
| Regulated on activation, normal T cell expressed and secreted/  C-C Motif Chemokine Ligand 5 | RANTES/  CCL5 | |
| Vascular Endothelial Growth Factor | VEGF | |

**Table S2. Representative peak shifts of control and irradiated MFPs detected by Raman spectroscopy.** Spectra were normalized to the 1440 cm^-1^ biological peak.

| Raman Shift (cm^-1^) | Assignment | Control | Irradiated |
| --- | --- | --- | --- |
| 1440 | Protein/Lipid | 1 | 1 |
| 1330 | Lipid | 0.475 | 0.453 |
| 1265 | Amide III (collagen) | 0.409 | 0.516 |
| 1660 | Amide I (collagen) | 0.537 | 0.731 |
| 1070 | DNA | 0.385 | 0.383 |
| 927 | Protein | 0.204 | 0.316 |
| 1126 | Lipid/Protein/Proteoglycan | 0.282 | 0.368 |
